# Parkinson’s disease-related Miro1 mutation induces mitochondrial dysfunction and loss of dopaminergic neurons *in vitro* and *in vivo*

**DOI:** 10.1101/2023.12.19.571978

**Authors:** Axel Chemla, Giuseppe Arena, Ginevra Sacripanti, Kyriaki Barmpa, Alise Zagare, Pierre Garcia, Paul Antony, Jochen Ohnmacht, Alexandre Baron, Jaqueline Jung, Anne-Marie Marzesco, Manuel Buttini, Thorsten Schmidt, Anne Grünewald, Jens C. Schwamborn, Rejko Krüger, Claudia Saraiva

## Abstract

The complex and heterogeneous nature of Parkinson’s disease (PD) is still not fully understood, however, increasing evidence supports mitochondrial impairments as a major driver of neurodegeneration in PD. Recently, the regulator of mitochondrial homeostasis Miro1 has been linked genetically and pathophysiologically to PD. Using 2D and 3D patient-based induced pluripotent stem cells models, including an isogenic control, showed that the Miro1 p.R272Q mutation leads to mitochondrial impairments including increased oxidative stress, disrupted mitochondrial bioenergetics and altered metabolism. This was accompanied by increased α-synuclein levels in 2D dopaminergic neurons and by a significant reduction of dopaminergic neurons within midbrain organoids. Knock-in mice expressing mutant p.R285Q Miro1 (orthologue of the human p.R272Q mutation) confirmed the PD-specific dopaminergic neuronal loss in the substantia nigra, accumulation of striatal phosphorylated α-synuclein accompanied by behavioral alterations. These findings demonstrate that mutant Miro1 is sufficient to comprehensively model PD-relevant phenotypes *in vitro* and *in vivo*, reinforcing its pivotal role in PD pathogenesis.

## Introduction

Parkinson’s disease (PD) is the fastest growing neurodegenerative disorder worldwide [1]. The majority of cases are sporadic, while around 5-15% are familial, caused by rare, high penetrance mutations in single genes showing autosomal dominant (e.g. *SNCA*, *LRRK2*) or recessive (e.g. *PINK1*, *PRKN*, *DJ-1*) inheritance [2]. Studies on these rare monogenic PD forms provided important insight into cellular and molecular mechanisms underlying neurodegeneration, such as mitochondrial dysfunction, oxidative stress, calcium dyshomeostasis, impaired autophagy and mitophagy, protein misfolding and apoptosis, among others [3]. These could cause the massive loss of dopaminergic neurons observed in the substantia nigra pars compacta (SNpc) of the PD brain, which prompts the well-described motor and some of the non-motor symptoms typical of PD. Surviving neurons in affected brain regions usually display characteristic proteinaceous aggregates (Lewy bodies) containing α-synuclein as a major component [3].

In sporadic PD patients, besides the impact of common genetic variants explored in large genome-wide association studies, the identification of disease-associated rare genetic variants is increasing, suggesting the contribution of low-frequency variants to disease susceptibility [4]. We previously identified four PD patients carrying distinct heterozygous mutations in the *RHOT1* gene encoding Miro1 [5,6]. Miro1 (mitochondrial Rho GTPase protein) is a conserved element of the mitochondria motor/adaptor complex. It is composed of a C-terminal transmembrane domain, which binds to the outer mitochondrial membrane, and two EF-hand Ca^2+^-binding domains flanked by two GTPase domains [7]. Miro1 plays a fundamental role in regulating not only mitochondrial dynamics (including morphology and transport), but also calcium homeostasis and mitophagy [8]. Miro1 was also found to physically or functionally interact with well-established PD-related proteins, such as PINK1, Parkin, LRRK2 and α-synuclein, suggesting a possible converging role of Miro1 in controlling different cellular activities and pathways relevant for neurodegeneration in PD [9–13]. Moreover, a pathological stabilization of Miro1, in which its degradation induced by mitochondrial depolarization is impaired, was observed in fibroblasts [10] and induced pluripotent stem cells (iPSC) [14] from both, sporadic and monogenic, PD patients. In accordance, the pharmacological reduction of Miro1 rescued mitochondrial dysfunction and neurodegeneration in different PD models *in vitro* and *in vivo*, including human iPSC-derived neurons from affected *LRRK2*-G2019S, *SNCA*-A53T carriers and sporadic PD patients as well as Drosophila [9–11]. Moreover, in human-derived fibroblasts and iPSC-derived neurons mutations in *RHOT1* have showed mitochondria-related alterations [6,15–17].

In the current study, we dissected the importance of Miro1 in PD pathogenesis by using iPSC-derived dopaminergic neuronal cultures and 3D midbrain organoids (MO), including an isogenic control derived from a PD patient carrying the p.R272Q mutation in Miro1 together with sex and age-matched healthy individuals as controls. We demonstrated that p.R272Q Miro1 mutation lead to an increase in reactive oxygen species (ROS) and altered mitochondrial bioenergetics. This was accompanied by higher α-synuclein levels and loss of dopaminergic neurons *in vitro*. Importantly, degeneration of dopaminergic neurons and accumulation of phospho-α-synuclein was observed in the SNpc of aged knock-in mice expressing the orthologue p.R285Q mutant Miro1, accompanied by deficits in the first trial of the Rotarod behavior test. Altogether, our findings indicate that PD-associated mutant Miro1 is sufficient to model PD *in vitro* and *in vivo*, supporting a role of Miro1 in the pathogenesis of PD.

## Results

### Miro1 p.R272Q mutation in midbrain organoids and dopaminergic neurons revealed transcriptomic deregulation of major PD-related pathways

To dissect the relevance of Miro1 in PD we used two different iPSC-derived *in vitro* models. We generated midbrain organoids, 3D complex structures with diverse cell types, including functional dopaminergic neurons, and defined spatial orientations mimicking the human midbrain [18,19]. Our midbrain organoids have also proven to be capable to model Parkinson’s disease phenotypes in a robust manner [20,21]. Additionally, we used 2D dopaminergic neurons to dissect in more detail the effects on the main affected cellular population in PD. These models were generated from iPSC obtained from a PD patient carrying the *RHOT1* c.815G>A (NM_001033568; Miro1 p.R272Q, first EF-hand domain) mutation (PD-R272Q), its correspondent line with point mutation correction (isogenic control, iCtrl) [22] as well as control lines from age- and sex-matched healthy individuals (Ctrl) (Figure 1A; Table1).

**Figure 1.**
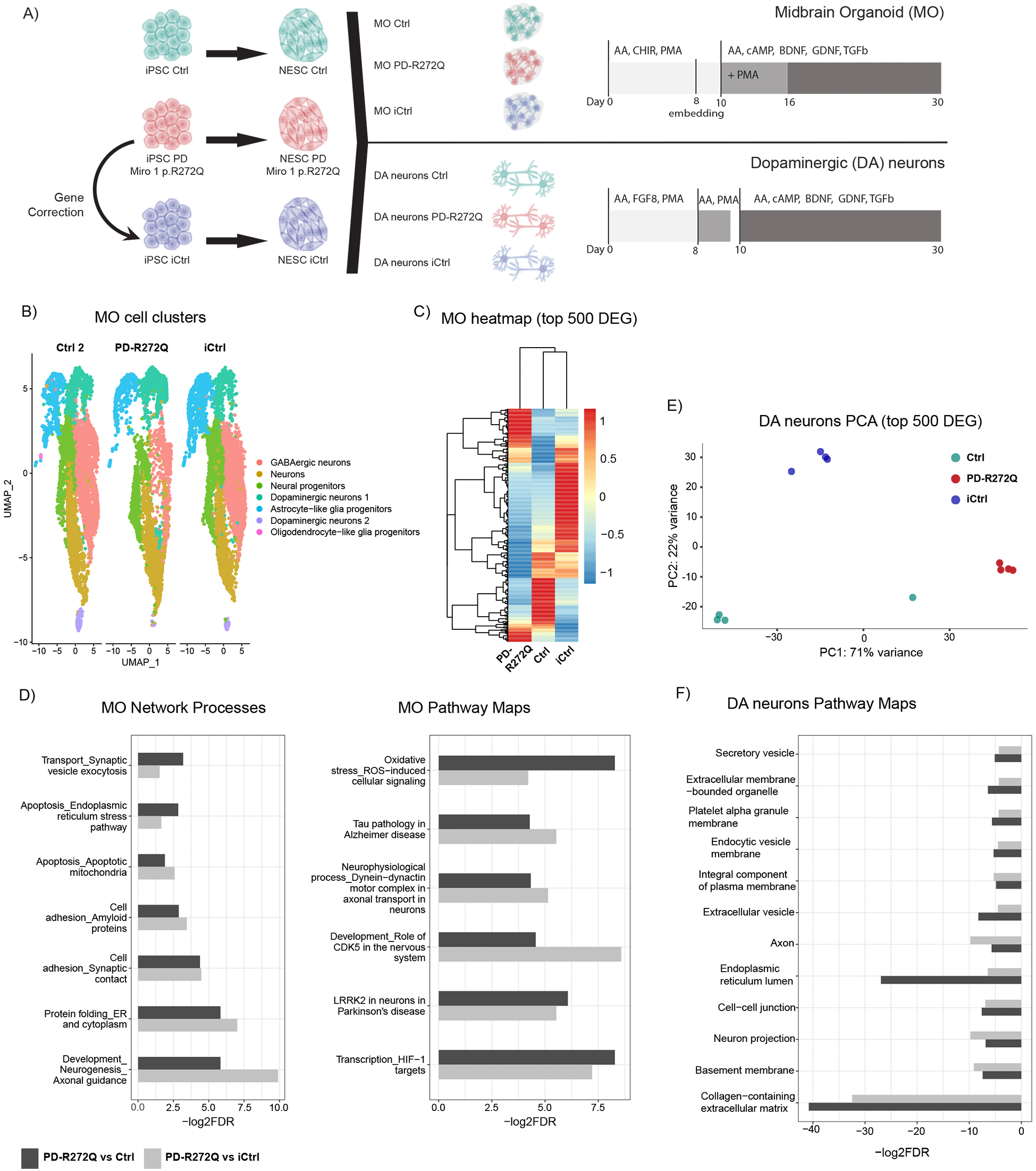
Parkinson’s disease (PD)-related pathways were deregulated in Miro1 p.R272Q mutant midbrain organoids (MO) and dopaminergic (DA) neurons. A) Schematic representation of the *in vitro* models and culture conditions used. B) UMAP visualization of healthy control (Ctrl 2), Miro1 p.R272Q mutant (PD-R272Q) and isogenic control (iCtrl) organoids single cell RNA sequencing (scRNAseq) data showed 7 unique cell clusters. Dots are color code by cell cluster and represent individual cells. C) Heatmap displaying the top 500 differential expressed genes (DEG) showed a separation between PD-R272Q MO and controls. D) Graphics depict the most PD-relevant deregulated network processes (left) and pathways (right) between MO PD-R272Q and Ctrl or iCtrl from the top 25 most deregulated ones (See also Supplementary Figure 2). E) Principal component analysis (PCA) plot showed separation at PC1 in PD-R272Q dopaminergic neurons compared with Ctrl and iCtrl, based on the top 500 DEG identified from bulk RNA sequencing analysis. F) Graphic displaying the deregulated gene ontology (GO) terms of PD-R272Q dopaminergic neurons *versus* healthy or isogenic controls. For all, significance was considered when p.adjusted value < 0.05.

**Table 1.** Description of cell lines used in *in vitro* experiments, both for midbrain organoids and dopaminergic neuronal cultures.

| ID | Diagnosis | Genotype | Sex | Age of sampling | Age of onset | Reference/Source |
| --- | --- | --- | --- | --- | --- | --- |
| Ctrl1 | Healthy (Control) | wt/wt | Female | 72 | - | [15] |
| Ctrl2 |  | wt/wt | Female | 68 |  | [52] |
| Ctrl3 |  | wt/wt | Female | 63 | - | [52] |
| PD-R272Q | PD | Miro1<br>p.R272Q/wt | Female | 78 | 70 | [22] |
| iCtrl | PD background /<br>Healthy genotype<br>(Isogenic control) | wt/wt | Female | 78 | 70 | [22] |

Midbrain organoids derived from 1 healthy individual (Ctrl2), PD-R272Q and iCtrl were analyzed using single cell RNA sequencing (scRNAseq). The Seurat integration workflow identified 7 different cellular clusters (Figure 1B) based on the combination of the La Manno et al. gene list [23] and the expression of cell and maturity specific markers (Supplementary Figure 1A-B). UMAP representation showed the presence of midbrain relevant cell types in the organoid model, including the presence of a neural progenitor cluster as well as two dopaminergic neuron clusters: dopaminergic neurons 1 and dopaminergic neurons 2, the later expressing higher levels of tyrosine hydroxylase (TH; Figure 1B, Supplementary Figure 1B). Hierarchical clustering of the top 500 variable genes showed that PD-R272Q organoids were separated from both healthy and isogenic controls, suggesting a significant influence of the Miro1 p.R272Q mutation on the transcriptome (Figure 1C). Then, the computed differentially expressed genes (DEG) were used to perform pathway enrichment analysis between PD-R272Q and either Ctrl or iCtrl midbrain organoids. The most relevant shared dysregulated processes and pathways (Figure 1D) from the top 25 (Supplementary Figure 2A) showed a significant deregulation of processes related to neurogenesis, synapses contacts and exocytosis, endoplasmic reticulum (ER) and mitochondrial apoptosis. Pathway maps also showed deregulation of ROS, transcription of HIF-1 targets (important for iron homeostasis and oxidative stress defense [24]), LRRK2 in PD neurons, and dynein-dynactin motor complex in axonal transport in neurons. Remarkably, the main transcriptomic alterations induced by mutant Miro1 seemed to be driven by the dopaminergic neuron clusters (Supplementary Figure 2B).

The transcriptomic profile of iPSC-derived dopaminergic neurons was also assessed using bulk RNAseq. Principal component analysis (PCA) showed a clear separation between PD-R272Q and controls (Figure 1E; Supplementary Figure 3). Gene ontology (GO) and enrich terms analyzed in PD-R272Q vs Ctrl and PD-R272Q vs iCtrl conditions showed deregulation in neuronal projections, axons, secretory vesicles, ER and lysosomal compartments (Figure 1F). These results further support specific Miro1 p.R272Q-dependent transcriptome alterations within dopaminergic neurons.

### Miro1 p.R272Q mutant midbrain organoids and dopaminergic neurons showed signs of mitochondrial stress

Pathway enrichment analysis on midbrain organoids scRNAseq data revealed several deregulated genes associated with ROS production (Figure 2A, Supplementary Figure 4), including: i) *ADM*, which is also involved in neurotransmitter release and cell adhesion [25]; ii) *FTL*, essential for iron metabolism and iron-induced stress [24]; iii) *PRKCB*, encoding protein kinase C beta type, which negatively correlates with mitochondrial energetic state and autophagy [26]; iv) *MAPK8* and *MAPK10*, mitogen-activated protein kinases, involved in misfolded protein-induced stress, autophagy, protein transport and apoptosis [27]; and v) *UBL5*, important for coping with mitochondrial stress [28]. Deregulation of these ROS-related genes pointed to a possible impairment of mitochondrial homeostasis in PD-R272Q organoids. Western blotting against the mitochondrial outer membrane protein VDAC (voltage-dependent anion channel) revealed that mitochondrial mass in organoids was similar between PD-R272Q and both controls (Figure 2B). MitoSOX Red was used to measure mitochondrial superoxide levels via flow cytometry. Mutant Miro1 p.R272Q organoids showed a significant increase in the number of MitoSOX-positive events compared with healthy and isogenic controls (Figure 2C). Mitochondria membrane potential (MMP) was assessed by the percentage and the mean fluorescent intensity (MFI) of the TMRM-positive signal within the total mitochondria (MitoTracker Green signal). Midbrain organoids expressing mutant p.R272Q Miro1 showed a significant reduction of the percentage of mitochondria with intact MMP as well as an overall lower MMP in comparison to controls (Figure 2D). Midbrain organoids from the iCtrl showed a significantly higher MMP compared to healthy controls.

**Figure 2.**
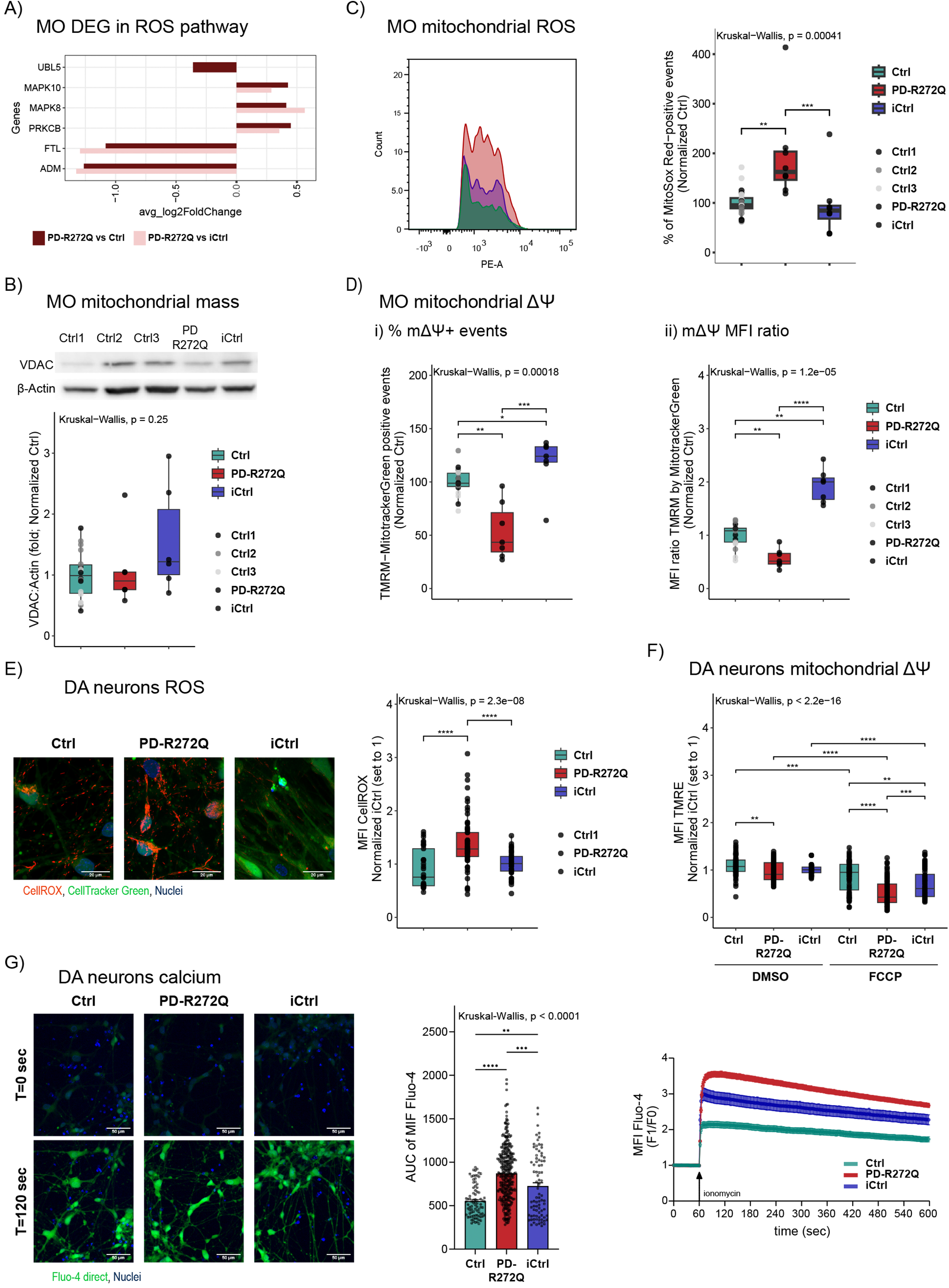
Miro1 p.R272Q mutation increased ROS, impaired mitochondrial membrane potential (MMP) and calcium handling *in vitro*. **A)** Fold change of selected significantly differentially expressed genes between Miro1 p.R272Q midbrain organoids (PD-R272Q MO) and healthy (Ctrl) or isogenic (iCtrl) controls, obtained from the enriched deregulated pathway *‘oxidative stress ROS-induced cellular signaling’* (see also Supplementary Figure 4). **B)** Box plot (bottom) showing MO Western blotting against the outer mitochondrial membrane protein VDAC (32 kDa) normalized to the housekeeping protein β-actin (42 kDa). n = 6-18 from 6 independent derivations. Representative images displayed on top. **C)** Flow cytometry representation (left) and quantification (right) of the percentage of MitoSox Red-positive events. n = 8-24 from 6 independent derivations. **D)** Evaluation of MMP using the specific marker TMRM by flow cytometry in organoids. **(i)** Percentage of double positive events for TMRM and Mito tracker Green. **(ii)** Mean fluorescent intensity (MFI) ratio between TMRM and Mito tracker Green within the double positive events. n = 5-21 from 5 independent derivations. **E)** Evaluation of cellular ROS in dopaminergic neurons using live imaging. Left: representative images of dopaminergic neurons containing CellRox deep red (ROS marker; red), Cell tracker green (cellular marker; green) and Hoechst (blue, nuclei) in Ctrl, PD-R272Q and iCtrl. Scalebar: 20 µm. Right: graphs depicting the intracellular CellRox MFI values normalized to iCtrl. n = 33-68, from 5 independent differentiations. **F)** MMP evaluation in dopaminergic neurons, in absence (DMSO) or presence of FCCP by live imaging quantification of mitochondrial TMRE MFI signal (boxplot). n = 63-84 from 4 independent differentiations. **G)** Left: representative images of dopaminergic neurons containing Fluo-4 direct (green), and Hoechst (blue, nuclei) before (zero seconds) and after (120 seconds) ionomycin injection. Scalebar: 50 µm. Middle: Bar plot shows F1/F0 area under the curve (AUC; right). Right: Graphical representation of Fluo-4 direct signal (F1) divided by its basal signal (F0) throughout time assessed by live imaging. n = 84 Ctrl1, n = 344 PD-R272Q, n = 82 iCtrl from 4 independent differentiations (left). All data is presented as median with max/min or mean ± SEM, *P < 0.05, **P < 0.01, ***P < 0.001 using non-parametric multiple comparison Kruskal-Wallis test.

In order to understand which of these phenotypes were specific for dopaminergic neurons, we characterized Miro1 p.R272Q mitochondria using dopaminergic neuronal cultures. Imaging-based analyses were performed to assess intracellular ROS, MMP and calcium response. Intracellular ROS were quantified by the MFI of the CellROX Deep Red-positive cells from total cells (Cell Tracker Green-positive signal). In concordance with midbrain organoid data, PD-R272Q neurons showed a significant increase of ROS compared to controls (Figure 2E). For MMP measures, MFI of TMRE (marker of intact MMP) within the total mitochondrial signal was quantified both in basal conditions (DMSO, dimethyl sulfoxide) and under mitochondrial depolarization (FCCP). In DMSO-treated cells, MMP was significantly lower in PD-R272Q neurons compared to the Ctrl, but not to the iCtrl. However, in response to mitochondrial depolarization, the Miro1 p.R272Q mutant neurons showed a reduced ability to cope with stress, and displayed a significantly reduced MMP compared with both controls (Figure 2F). Finally, due to the nature of the Miro1 p.R272Q mutation that affects Miro1 EF-calcium binding domain [6,15,16], and the existing link between impaired cytosolic calcium handling and mitochondrial stress [29], calcium response was evaluated. Fluo-4 Direct MFI was measured over time in presence of ionomycin, which promotes a rapid influx of Ca^2+^ into cells. PD-R272Q dopaminergic neurons showed the highest increase in the F1/F0 ratio upon ionomycin administration (Figure 2G). Herein, iCtrl dopaminergic neurons showed an intermediate response to the increased intracellular calcium levels, displaying significant differences compared to both Ctrl and PD-R272Q neurons (Figure 2G). Altogether, our results showed that despite some influence of the patient’s genetic background, the Miro1 p.R272Q mutation seems to increase the susceptibility to mitochondrial damage, which might impact mitochondrial energy production.

### Miro1 p.R272Q-induced mitochondria stress prompted bioenergetic deficits *in vitro*

To understand the influence of the Miro1 p.R272Q mutation on the energy status of the cells, we analyzed mitochondrial respiration in midbrain organoids using the Seahorse technology. As shown in Figure 3A (left panel), the pattern of oxygen consumption rate (OCR) curves was significantly different between PD-R272Q organoids and both, healthy and isogenic controls. In particular, PD-R272Q organoids showed a significantly lower basal respiration, proton leak and ATP-linked production. Moreover, PD-R272Q organoids displayed a significantly lower non-mitochondrial respiration compared to Ctrl but not to the iCtrl, suggesting a partial contribution of the patient’s genetic background. However, when treated with the mitochondrial uncoupler FCCP, PD-R272Q midbrain organoids were still able to respond in a similar way as control organoids, shown by the similar levels of maximal respiration and higher spare respiratory capacity observed (Figure 3A, right panel). Midbrain organoids metabolomic analysis did not show a significant difference in ATP levels (Figure 3B i), indicating potentially compensatory mechanisms from non-mitochondrial metabolic pathways. Nevertheless, PD-R272Q organoids showed significant lower relative abundance of the co-factors NAD and FAD (Figure 3B ii, iii) as well as pyruvate, a key metabolite in the glucose metabolism that feeds the tricarboxylic acid (TCA) cycle (Figure 3B iv). We further assessed the specific role of the p.R272Q mutation in 2D human dopaminergic neurons, which showed a significant decrease of all OCR-related parameters compared to healthy and isogenic controls (Figure 3C). Interestingly, different from Miro1 p.R272Q organoids, dopaminergic neurons also displayed a significant reduction of maximal respiration and spare respiratory capacity (Figure 3C, right panel), indicative of their inability to produce energy via oxidative phosphorylation under high energy demand. The mitochondrial phenotype observed in PD-R272Q dopaminergic neurons was also supported by the robust decrease of intracellular ATP levels in comparison with both controls (Figure 3D). Levels of NAD(H) (Figure 3E i) and NADP(H) co-factors (Figure 3E ii) were also affected in PD-R272Q neurons when compared to isogenic and/or healthy controls, although no significant alterations were observed in their ratio. Interestingly, these energy alterations seemed to be more pronounced between PD-R272Q vs iCtrl than PD-R272Q vs Ctrl, indicating a Miro1 p.R272Q mutation-dependent effect. Metabolomic analysis of dopaminergic neurons media revealed that PD-R272Q neurons released more lactate (Figure 3F i) while taking more pyruvate (Figure 3F ii), glutamate (Figure 3F iii) and glycine (Figure 3F iv) up than iCtrl. Surprisingly, no significant differences were observed in the relative abundance of these metabolites between Ctrl and PD-R272Q neurons. Overall, these results support mitochondria-related bioenergetic defects caused by the p.R272Q Miro1 mutation.

**Figure 3.**
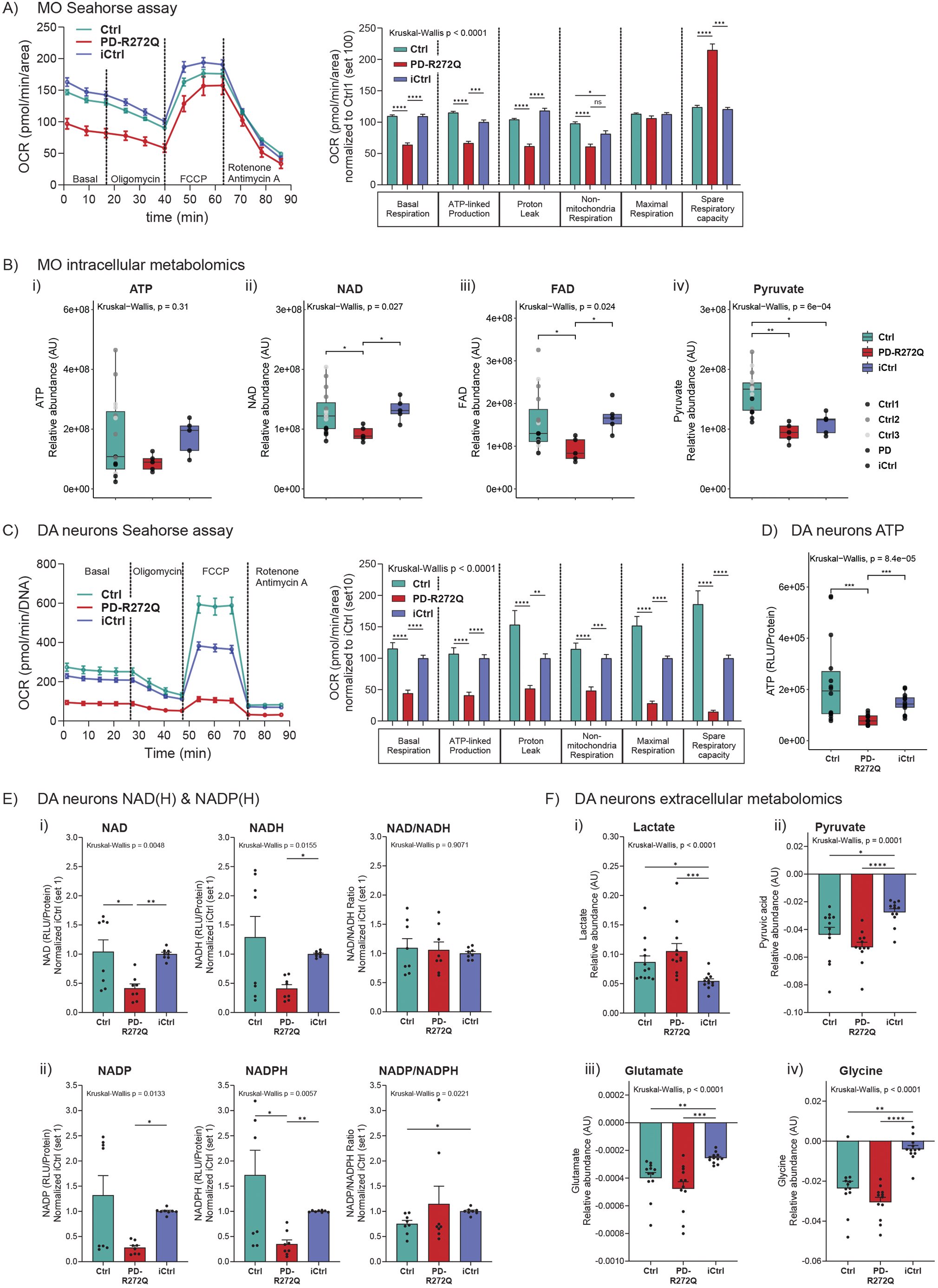
Miro1 p.R272Q mutation caused mitochondrial-based bioenergetic deficits *in vitro*. **A)** Assessment of Miro1 p.R272Q mutant (PD-R272Q), healthy (Ctrl) and isogenic (iCtrl) midbrain organoids (MO) mitochondrial function using Seahorse mito stress test from Agilent. Representative graphic depicts oxygen consumption rate (OCR) over time (minutes, min) under a specific set of drugs: oligomycin, FCCP and antimycin and rotenone. On the right, bar graph shows quantification of the different parameters calculated from the assay. Ctrl n = 197, PD n = 53, iCtrl n = 35 organoids from 5 (iCtrl) or 9 (Ctrl, PD-R272Q) independent derivations. **B)** Relative abundance of (i) ATP, (ii) NAD, (iii) FAD and (iv) pyruvate intracellular metabolites in MO upon targeted liquid chromatography – mass spectrometry (LC-MS). n = 5-15 from 2 independent derivations. **C)** Graphics display OCR by time (left) and feature quantification (right) from Seahorse Mito stress test in dopaminergic neuronal cultures; n = 29-34 from 3 independent differentiations. **D)** Intracellular ATP levels in dopaminergic neurons. n = 14 from 5 independent differentiations. **E)** Intracellular NAD(H) and ratio **(i)** as well as NADP(H) and their ratio **(ii)** relative quantification in dopaminergic neurons. n = 12 from 4 independent differentiations. **F)** Relative quantification of polar metabolites in dopaminergic neuron extracellular media using gas chromatography (GC)-MS. Graphs showed release of **(i)** lactate and uptake of **(ii)** pyruvate, **(iii)** glutamate, **(iv)** glycine metabolites. All data is represented as mean ± SEM or median with max/min. *P < 0.05, **P < 0.01, ***P < 0.001, ****P < 0.0001 using non-parametric multiple comparison Kruskal-Wallis test.

### Miro1 p.R272Q mutant dopaminergic neurons showed increased levels of total α-synuclein

Recent evidence points to a direct role of α-synuclein in mitochondrial dynamics and quality control mechanisms [30] as well as a possible positive correlation between α-synuclein and Miro1 levels [11]. In midbrain organoids, *RHOT1* gene expression levels were not different (Supplementary Figure 5A). Though, DEG analysis showed a significant increase in *SNCA* gene expression in PD-R272Q organoids compared with healthy and isogenic controls (Figure 4A, average expression levels: Ctrl = 0.35, PD-R272Q = 0.83, iCtrl = 0.35). Nevertheless, no difference in total α-synuclein protein levels were observed (Figure 4B). In 2D dopaminergic neurons, *RHOT1* was upregulated in PD-R272Q compared to Ctrl and iCtrl within the bulk RNAseq data (log2 fold change = 0.52 and 1.92, respectively; Supplementary Figure 5B). Likewise, *SNCA* transcript was upregulated in PD-R272Q compared to Ctrl and iCtrl (log2 fold change = 0.79 and 2.31, respectively) – (Figure 4C). In accordance, α-synuclein levels were significantly higher in PD-R272Q dopaminergic neurons compared to iCtrl but not to Ctrl (Figure 4D). These findings suggest that Miro1 p.R272Q might interfere with α-synuclein expression levels.

**Figure 4.**
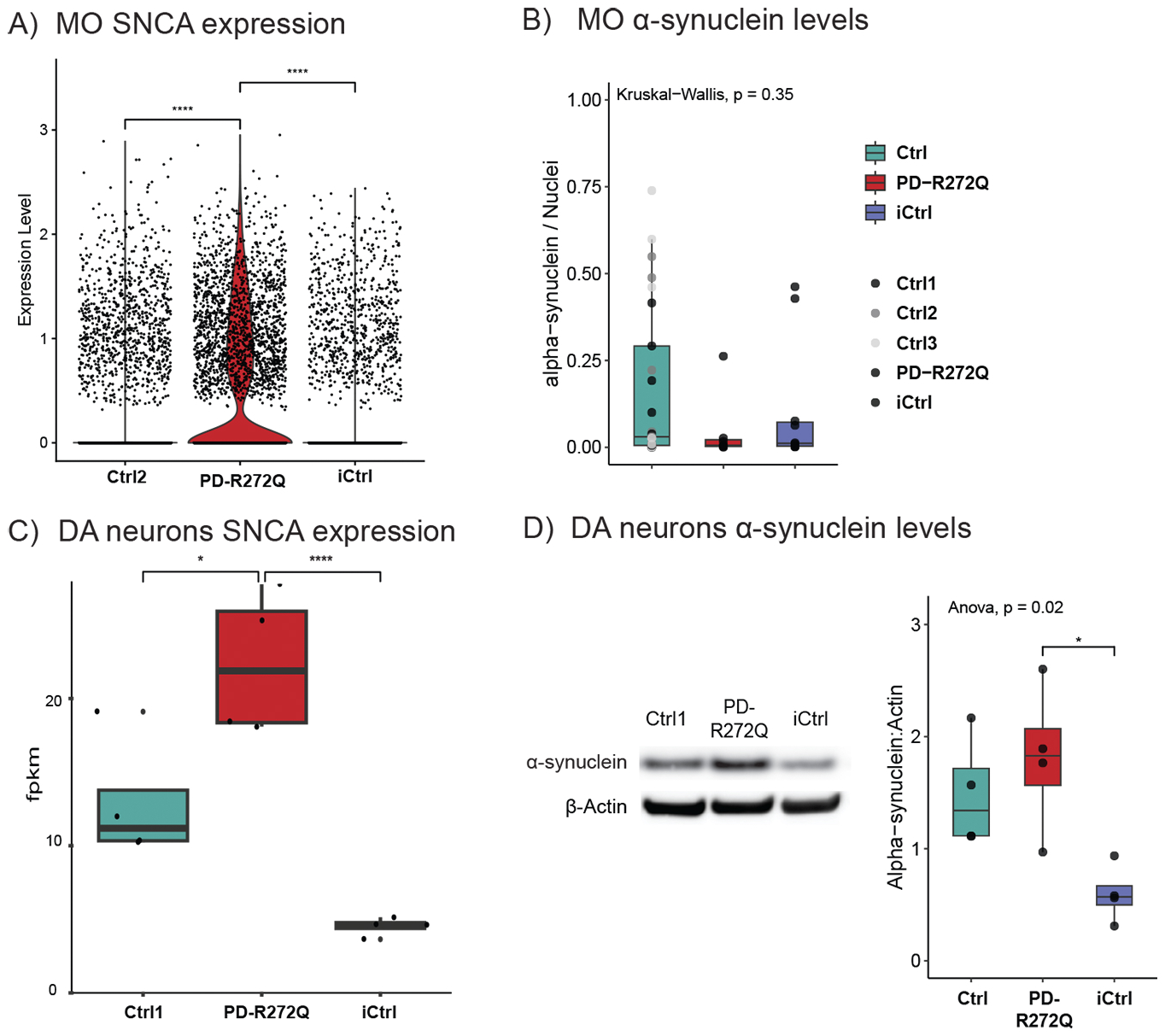
Dopaminergic neurons with Miro1 p.R272Q mutation showed higher α-synuclein levels. **A)** SNCA expression levels in midbrain organoids (MO) obtained from transcriptomic data. **B)** MO showed similar levels of α-synuclein in immunofluorescent staining. n =9-27 from 5 independent derivations. **C)** SNCA expression levels in dopaminergic neurons infer from bulk RNA sequencing analysis. **D)** Top: Representative image of α-synuclein (15 kDa) and housekeeping gene β-actin (42 kDa) Western blotting in dopaminergic neurons. Bottom: α-synuclein levels quantification. n = 4. Data presented as median with max/min. *P < 0.05, **P < 0.01, using non-parametric multiple comparison Kruskal-Wallis test **(B)** or ANOVA with post-hoc Tukey HSD test **(D)**.

### Increased dopaminergic neuronal cell death in Miro1 p.R272Q midbrain organoids

The loss of dopaminergic neurons is a key hallmark of PD. Immunoblotting analyses showed a significant reduction of the dopaminergic marker TH in PD-R272Q organoids compared with both healthy and isogenic controls (Figure 5A). Similar results were obtained by confocal microscopy, upon quantification of the TH-positive (TH+) signal within the total neuronal population (TUJ1+) (Figure 5B). High content image morphometric analysis [31,32] revealed that, in mutant organoids, TH+ neurons have a reduced average length (skeleton) compared to Ctrl, but not to iCtrl (Figure 5C i), meaning that TH+ PD-R272Q neurons have in average a smaller cytoskeleton. Moreover, the neurite fragmentation index of TH+ neurons, which is an early indicator of neurodegeneration [33], was significantly increased in PD-R272Q organoids in comparison with both Ctrl and iCtrl (Figure 5C ii). In line, scRNAseq analysis showed a clear deregulation of genes related with apoptosis within the dopaminergic neuron clusters (Supplementary Figure 6). Lactate dehydrogenase (LDH) presence in organoids media and the TUNEL apoptotic assay were used to assess cell death. PD-R272Q organoids showed a significant increase in LDH abundance (Figure 5D) and higher levels of DNA fragmentation (as measured by the TUNEL staining; Figure 5E) within the dopaminergic neuron population, compared to Ctrl and iCtrl. Altogether, these findings suggest a selective loss of TH+ neurons in Miro1 mutant organoids, which is mediated, at least in part by apoptosis.

**Figure 5.**
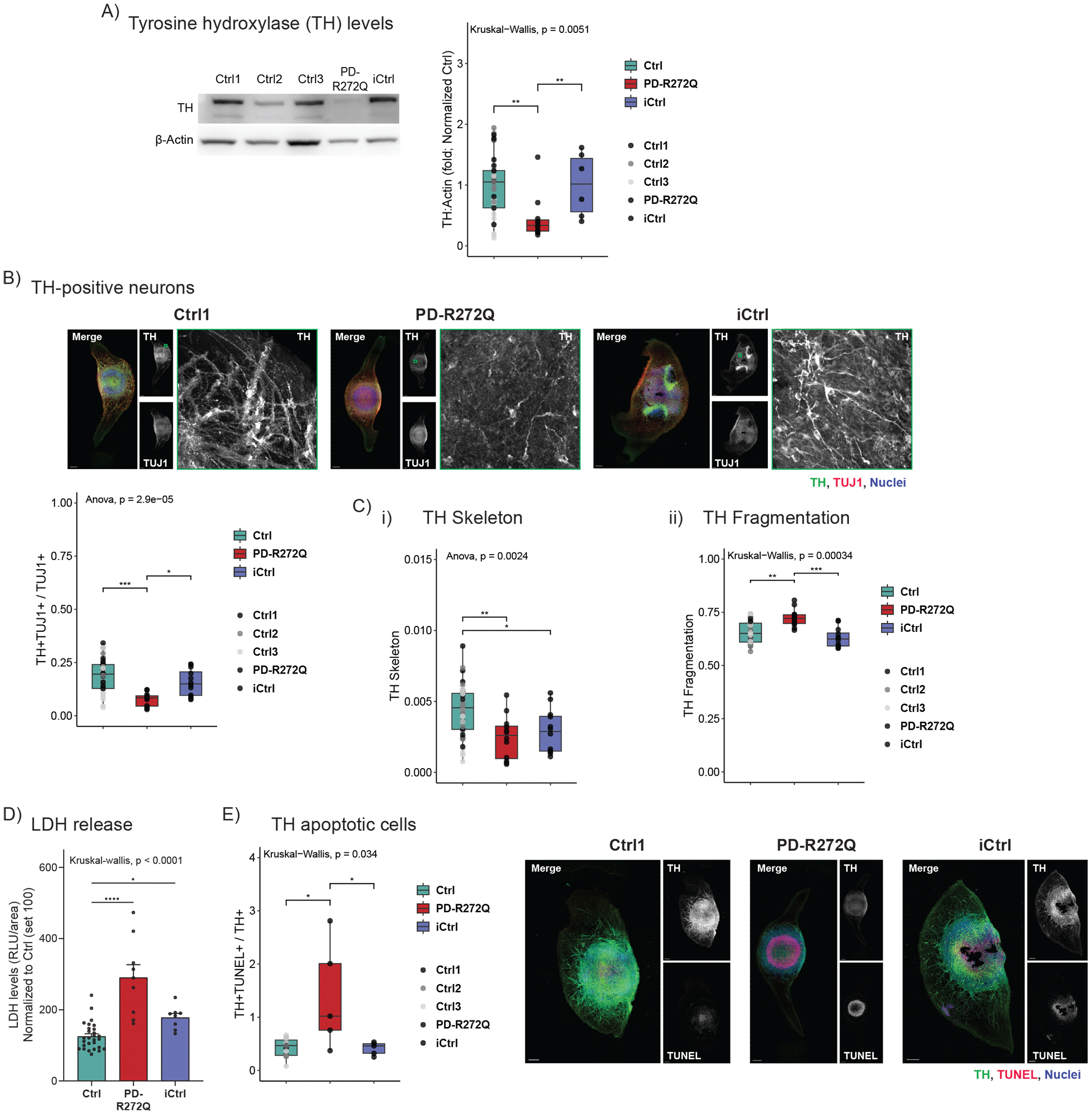
Miro1 p.R272Q mutant midbrain organoids (MO) showed signs of dopaminergic neuron loss. **A)** Tyrosine hydroxylase (TH) quantification (right) and representative image (left) evaluated in organoids by Western blotting (TH – 60 kDa; β-actin – 42 kDa). Ctrl n = 30, PD-R272Q n = 10, iCtrl n = 6 from 6 (iCtrl) or 10 (Ctrl, PD) independent derivations. **B)** Immunofluorescent analysis of TH-positive (TH+) signal, total neural marker TUJ1 and nuclei (Hoechst) in MO. TH area Scale bar: 200 µm. Graphic displays quantification of the total volume occupied by the TH and TUJ1 double positive signal within the total TUJ1. **C)** Immunofluorescent TH-based morphometric features **(i)** 3D Skeleton and **(ii)** fragmentation index. **B, C)** n = 12-36 from 5 independent derivations. **D)** Quantification of the relative abundance of lactate dehydrogenase (LDH) released into MO media. n = 8-28 from 3 independent derivations. **F)** Immunofluorescent identified TH+ neurons undergoing apoptosis (TUNEL assay). Nuclei are shown in blue. Scale bar: 200 µm. Box plot depicts volume of TH neurons undergoing apoptosis (TH+TUNEL+) normalized by the total TH. n = 7-16 organoids from 3 independent derivation. All data is presented as median with max/min. or mean ± SEM. *P < 0.05, **P < 0.01, ***P < 0.001, ****P < 0.0001 using non-parametric multiple comparison Kruskal-Wallis test **(A, Cii, D, E)** or ANOVA with post-hoc Tukey HSD test **(B, Ci)**.

### Aged Miro1 p.R285Q knock-in mice showed dopaminergic neuron degeneration and behavioral alterations

The Miro1 protein is evolutionary conserved among species [8]. The R285 amino acid in mice is the equivalent of the human R272 residue found mutated (i.e., p.R272Q) in the PD patient. Thus, to study the potential contribution of mutant Miro1 to PD pathogenesis *in vivo*, we created a knock-in mouse model by introducing the corresponding Miro1 p.R285Q mutation using CRISPR/Cas9 technology (Figure 6A). Successful generation of knock-in mice was validated by PCR and DNA sequencing analysis in wild-type animals (wt/wt) as well as in heterozygous (wt/R285Q) and homozygous (R285Q/ R285Q) mutant mice (Figure 6A). Miro1 mutation did not interfere with mice weight (Supplementary Figure 7A). Immunohistochemical analysis showed similar levels of TH-positive terminals and dopamine transporter (DAT) in the striatum (Supplementary Figure 7 B,C) and similar TH levels in the SNpc (Supplementary Figure 7D) of young mice (3-6 month-old, Figure 6B). In 15-month-old mice, striatal TH (Figure 6 C), DAT levels (Supplementary figure 7E) and striatal dopamine (Supplementary Figure 7F) were also unaffected. However, 15-month-old mutant mice showed a significant decrease in the SNpc TH area, reflecting dopaminergic neuron loss in both heterozygous and homozygous mice, an effect that was more pronounced in females (Figure 6D and Supplementary figure 7G). Considering this and the lower number of male mice within the study we proceed the analysis focusing on female mice. Total levels of α-synuclein in the striatum of old mice did not differ (Figure 6E). Nevertheless, old homozygous mutant mice showed significantly more phosphorylated S129 α-synuclein in the striatum (Figure 6F). The presence of phosphorylated S129 α-synuclein inclusions were also qualitatively observed via immunofluorescence in the mice SNpc (Figure 6G and Supplementary figure 7H). 20 month-old mice were monitored for water and food consumption as well as global activity using Phenomaster® cages. No significant differences were observed between the mice (Supplementary Figure 7I, J, K). Then, Rotarod test was used to evaluate spontaneous motor activity [34] and/or anterograde procedural memory [35] in 20 (trial 1) and 21 (trial 2) month-old female mice. Heterozygous and homozygous Miro1 p.R285Q mice displayed a significantly reduced latency to fall compared to wild-type mice on their first trial (Figure 6H), but not in the second trial (Figure 6I), which is indicative of compromised motor or motor-learning processes. Taken together, characterization of Miro1 mutant mice suggests a pathological role of p.R285Q (human p.R272Q) point mutation in PD etiopathogenesis.

**Figure 6.**
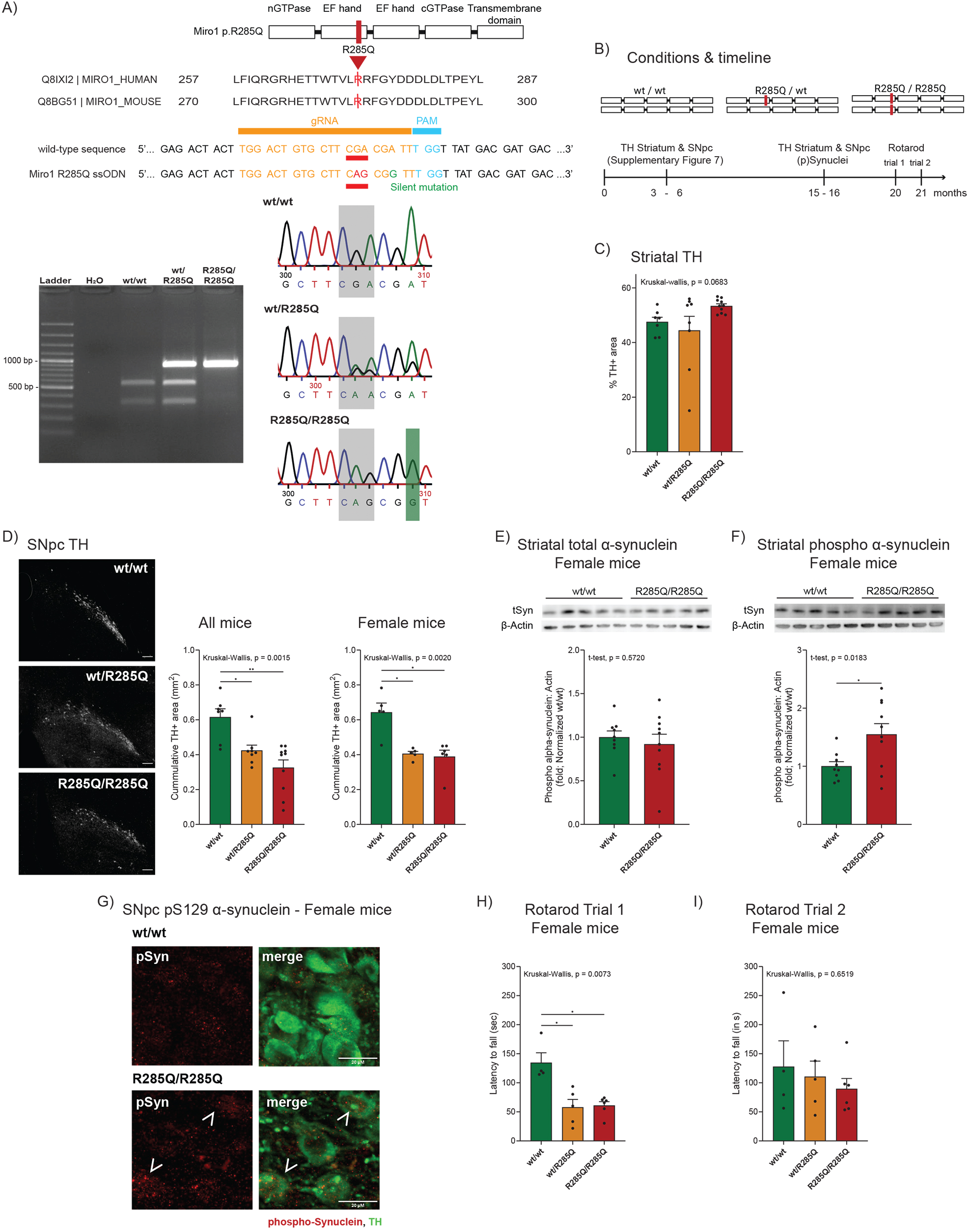
Miro 1 p.R285Q mutant aged mice presented dopaminergic neuron loss and behavior alterations. **A)** Protein amino acid sequencing and CRISPR/Cas9 guided RNA sequences used in the knock-in Miro1 p.R285Q mouse generation. Agarose gel (left) and DNA sequencing (right) confirming the successful integration of the Miro1 point mutation. **B)** Scheme illustrating the timeline of the experiments and mice used within the study: wildtype mice (wt/wt), heterozygous (wt/R285Q) and homozygous (R285Q/R285Q). **C)** Graphic depicts percentage of striatal tyrosine hydroxylase-positive (TH+) area. **D)** Representative image of SNpc TH immunoreactivity. Scale bar: 50µm. Quantification of the area occupied by TH in substantia nigra par compacta (SNpc) in 15 month-old mice (left) or in female only (right). **C, D)** wt/wt: n = 2M + 5F; wt/R285Q: n = 3M + 5F; R285/R285Q: n = 4M + 6F. **E, F)** Western blotting quantification (bottom) and representative image (top) of total α-synuclein **(E)** and phosphorylated (p)S129 α-synuclein **(F)** in the striatum of 15 old-month female mice. wt/wt n = 9, R285Q/R285Q n = 10. **G)** Representative confocal images of pS129 α-synuclein (red) and TH-positive (green) signal in the SNpc of 15 month-old female wildtype (top) and homozygous (bottom) mice. Arrows depicts pS129 α-synuclein intracellular inclusions. Scalebar 20µm. **H, I)** Bar graph display 21 month-old female mice latency to fall in seconds upon a first **(H)** and second **(I)** trials of Rotarod behavior test. wt/wt: n = 4; wt/R285Q: n = 5; R285/R285Q: n = 6. Graphs are represented as mean ± SEM, each point represents one mouse. *P < 0.05, **P < 0.01 using non-parametric multiple comparison Kruskal-Wallis test **(C, G, H)** or unpaired T test **(D, E)**. Abbreviation: F, female; M, male.

## Discussion

PD is not only a clinically but also genetically heterogenous disease. Besides rare Mendelian mutations in monogenic forms and common variants in genes associated with the common sporadic form of PD, there is growing evidence pointing to an important role of low-frequency variants with substantial effect in PD pathogenesis [36]. Recently, a large study investigating the burden of rare genetic variants in PD based on their predicted functional impact, identified *RHOT1* as one of four genes with the highest burden (4^th^ out 2500 genes) [37]. This strongly supports a role for rare variants in the *RHOT1* gene, encoding Miro1, including those previously identified by us [5,6]. Furthermore, a common molecular signature where Miro1 mitochondrial clearance is delayed (Miro1 retention phenotype) has been identified in both familial (*PINK1*, *PRKN*, *SNCA*, *LRRK2*) and sporadic PD cases [9–11], highlighting Miro1 as a potential convergent player, as well as a potential molecular biomarker in PD [38]. Due to limited pedigree size the genetic impact of *RHOT1* variants was not accessible to validation via classical Mendelian co-segregation patterns, urging the need for functional characterization. Herein, a novel mouse model in combination with patient-based iPSC-derived 2D and 3D cellular models, including a gene-corrected control, allowed for dissecting neuronal phenotypes related to the Miro1 p.R272Q mutation and their impact on PD pathogenesis. We demonstrated that Miro1 p.R272Q mutation causes mitochondrial dysfunction leading to dopaminergic neuron loss, both *in vitro* and *in vivo*. Importantly, we unveiled Miro1-dependent cellular processes and molecular signatures, providing convincing evidence to supporting its role in neurodegeneration in PD.

*In vitro*, transcriptomic analysis between Miro1 p.R272Q mutant and healthy or isogenic controls showed a mutation-specific deregulation on PD pathways, which are also known to be altered in sporadic and familial PD, such as LRRK2 and Tau deregulation, oxidative stress, iron homeostasis and apoptosis [39–41]. For example, the *ADM* gene encoding adrenomedullin – known to protect rat neurons against oxidative stress and apoptosis [42] – was downregulated in PD-R272Q organoids. Of note, the shorter pro-adrenomedullin fragment is implicated in increasing kinesin-mediated mitochondrial velocity [43], in line with our previous findings showing reduced mitochondrial speed in Miro1 p.R272Q dopaminergic neurons [15]. Other ROS-related deregulated genes, such as *UBL5* (ubiquitin-related) and *FTL* (iron-related) have been previously linked to Miro1-mediated mitophagy regulation [8] and to a putative iron-calcium-Miro axis [44], respectively. The Iron-calcium-Miro1 axis hypothesis states that elevated iron levels in PD lead to mitochondrial calcium overflow, which might be preceded by reduced calcium sensing Miro1 capability [44]. Indeed, despite the similar levels of total mitochondrial mass, in the PD-R272Q condition, we observed high levels of ROS and a reduced number of functional mitochondria. Miro1 is able to sense both cytosolic calcium [45] and oxidative stress [46,47], with impairments in Miro1 leading to mitophagy alterations and bioenergetics deficits [44,46,47]. A recent study in iPSC-derived dopaminergic neurons, from a healthy individual in which the heterozygous Miro1 p.R272Q mutation had been artificially introduced, showed Ca^2+^ deregulation as well as altered mitochondrial morphology and basal respiration [16]. In our patient-specific PD-R272Q dopaminergic neurons, we also showed calcium handling dysregulation [15], which might contribute to mitochondrial defects observed. Miro1 p.R272Q isogenic control showed an intermediate calcium response, further strengthen the concept that additional factors defining the individual ‘genetic background’ may influence the occurrence of particular phenotypes [48].

Mitochondrial respiration and energy production were severely impaired in both Miro1 mutant midbrain organoids and dopaminergic neurons (i.e., OCR, NAD and FAD abundance as well as metabolic alterations). These phenotypes were seen more clearly in the 2D dopaminergic neurons, which can be explained by the presence of other cellular populations within midbrain organoids. In fact, Miro1 p.R272Q mutant organoids showed a specific loss of TH-positive dopaminergic neurons in comparison with healthy and isogenic controls at day 30 of culture. The dopaminergic neuron population found within the midbrain organoids produces dopamine and has pacemaker activity [19]. These features can justify the higher vulnerability of the dopaminergic neuron population to stress and energy loss driven by the Miro1 p.R272Q mutation within the midbrain organoids, modeling the main hallmark of PD [3]. Likewise, loss of TH phenotype in the midbrain organoid model have also been demonstrated by us in familial forms of PD, including LRRK2, PINK1 and GBA mutations [20,21,49]. Furthermore, NAD(H) and NADP(H) co-factors were significantly reduced in PD-R272Q dopaminergic neurons, suggesting that energy deficits might come from low availability of these species – essential for cellular function – rather than their imbalance [50,51]. Supporting this, mitochondrial metabolic impairments and lower NAD pool have been reported in iPSC-derived neural precursors obtained from idiopathic PD patients [52]. Moreover, supplementation with NAD+ precursors niacin and nicotinamide riboside showed neuroprotective properties in PD patients [53,54].

Alpha-synuclein has recently been implicated in mechanisms controlling mitochondrial quality control and dynamics, such as autophagosome-lysosome formation and function, mitophagy, MDV formation, among others [30]. PD-R272Q dopaminergic neurons showed α-synuclein upregulation in a genetic background-dependent manner, suggesting a dopaminergic-specific α-synuclein accumulation. Previously, a positive correlation between Miro1 and α-synuclein levels was shown in the SNpc of sporadic PD patients [11], further supporting a possible functional link between both proteins. This link, might be related to altered mitochondrial quality control mechanisms [30] or Miro1-dependent calcium deregulation, since disruption in intracellular calcium buffering promotes α-synuclein aggregation [55], which can be caused by a direct interaction with the α-synuclein NAC domain [56]. Interestingly, transcriptomics in PD-R272Q organoids and dopaminergic neurons showed extracellular matrix-related genes dysregulation, which is involved in the pathological spreading of proteins (e.g. Tau, α-synuclein, beta-amyloid) typically observed in neurodegenerative disorders [57]. Altogether, our results further support a potential direct and/or indirect relation between Miro1 and α-synuclein dynamics, however further studies are needed to better understand the nature of their relationship.

*In vivo*, we generated the first knock-in mouse model of Miro1 p.R285Q mutant (ortholog of the human p.R272Q mutation) where physiological levels of Miro1 p.R285Q kept mimicking the human condition. Miro1 p.R285Q mutant mice presented significant dopaminergic neuronal loss in the SNpc of aged animals, corroborating the midbrain organoid experiments. Of note, these mice presented elevated levels of phosphorylated S129 α-synuclein, a post-translational modification that has been associated with α-synuclein toxicity and aggregation [58]. Indeed, reduction of S129 phosphorylated α-synuclein levels decreased motor deficits in PD mice overexpressing human α-synuclein [59]. Furthermore, p.R285Q mice spent significantly less time in the rod at the first trial of the Rotarod task, which might be due to difficulties in performing precise movement [34] or impaired anterograde procedural memory [35]. Dodson and colleagues showed that SNpc dopaminergic neurons stop firing at movement onset leading to slightly impairments in motor precision in humanized *SNCA* overexpression and *SNCA* knock-out mice models [60]. Procedural memory can be divided in the ability to learn new skills (anterograde), and the ability to remember and execute previously learned skills (retrograde). PD patients show decreased implicit learning abilities concerning new motor tasks (anterograde procedural memory), while their motor performance improves over time on pursuit rotor-motor task test [35,61]. Interestingly, rotarod differences in Miro1 p.R285Q mice were dissipated on trial two, suggesting either compensatory mechanisms or lack of pure motor deficits. Similarly, impairments on the first but not consecutive trials of the rotarod test were observed in a transgenic ataxia mouse model [62].

PD is commonly modeled by toxin-based acute challenges that, despite recapitulating SNpc neuronal loss, do not replicate the human PD pathological features [63]. In other PD mouse models, including single knock-out (e.g. *DJ-1, PINK1*, *PRKN*), triple *PINK1*/*PRKN*/*DJ-1* knock-out or *LRKK2*-R1441G transgenic mice, no dopaminergic neurodegeneration was observed albeit their motor coordination is altered [63–66]. Herein, we showed that physiological levels of Miro1 with p.R285Q mutation led to behavior impairments and SNpc TH loss in an age-dependent way. LRRK2 knock-in mice have also been used to model PD, showing subtle differences in striatal dopamine content and/or behavior defects [67]. However, dopaminergic degeneration was only observed in *LRKK2*-G2019S knock-in mice when combined with *SNCA*-A53T overexpression [68].In sum, these results point to the unequivocal key role of Miro1 in PD pathogenesis. Interestingly, our knock-in model had a more pronounced effect in females, which could be a sample size limitation and/or sex specific genetic or hormonal differences. Notable, the PD patient carrying the Miro1 p.R272Q is also female.

An important question still not addressed is how PD patient heterozygous Miro1 p.R272Q mutation affects Miro1 overall activity. The mutation is located in the calcium sensing domain, with a predicted toxic gain-of-function [6], supported by our current data. However, considering the current experimental setup and the Miro1 different domains further experiments are needed to clearly understand if p.R272Q is a gain-or loss-of function mutation. Miro1 knock-out is lethal, while conditional neuronal Miro1 knock-out or long deletion led to mitochondrial dysfunction and neurodegeneration [69,70]. However, Miro1 overexpression in animal fibroblasts also resulted in mitochondrial morphology and transport issues [71]. Similarly, we and others showed mitochondrial dysfunction in Miro1 mutant cells [6,15–17]. Remarkably, all Miro1 alterations culminate in similar mitochondrial dysfunction phenotypes, suggesting that a tight regulation of Miro1 levels and/or activity is needed for maintaining mitochondrial homeostasis, with slight changes being pathogenic for cells, especially for more sensitive cell types such as dopaminergic neurons. In fact, stabilization of wildtype Miro1 levels via downregulation [9–11,46] or overexpression [72] showed amelioration in mitochondrial-related phenotypes, relevant in PD.

## Conclusion

In conclusion, the here presented *in vitro* and *in vivo* models demonstrated that Miro1 p.R272Q mutation is sufficient to cause dopaminergic neuron loss, which might occur via alteration of mitochondrial status. We also showed a possible functional link between p.R272Q mutant Miro1 and α-synuclein. Moreover, we highlighted the importance of rare pathogenic variants, such as those found in *RHOT1*, as key in PD pathogenesis, further demonstrating the importance of genetic variance in PD susceptibility. Altogether, we corroborate the potential of Miro1 as convergent biomolecule and drug target in both familial and sporadic cases of PD as well as its relevance as a model of PD to develop new disease-modifying therapies.

## Competing interests

J.C.S. is co-inventor on a patent (WO2017060884A1) describing the midbrain organoid technology used and co-founder of OrganoTherapeutics (organoid-based biotech company). Other authors declare no competing interests.

## Funding

This work was mainly supported by the Luxembourg National Research Fund (FNR) under the CORE Junior Program (C19/BM/13535609; C.S. & J.C.S.) and the CORE grant MiRisk-PD (C17/BM/11676395; R.K., A.G. and G.A). R.K. also obtained funding from the FNR PEARL Excellence Program (FNR/P13/6682797), the Michael J. Fox Foundation, and the European Union’s Horizon2020 research and innovation program (WIDESPREAD; CENTRE-PD; grant agreement no. 692320). A.C. is supported by the FNR within the framework of the PARK-QC DTU (PRIDE17/12244779/PARK-QC). G.A. is also supported by the FNR CORE Junior grant PINK1-DiaPDs (C21/BM/15850547). A.G. was supported by the FNR within the ATTRACT program (Model-IPD, FNR9631103). The authors also thank Prof. Michel Mittelbronn funded by FNR PEARL (P16/BM/11192868).

For the purpose of open access, authors have applied a CC BY public copyright license to any author accepted manuscript (AAM) version arising from this submission.

## Author contribution

A.C.: Design of experiment, performed, analyzed and interpret 2D and mice immunohistochemistry experiments, manuscript writing and illustration. G.A.: Study design, conceptualization and management of mice work, data collection and interpretation, supervision of work, manuscript revision. G.S.: collection of 3D data. K.B. & A.Z.: analysis and plotting of organoid scRNAseq. P.A.: MATLAB script development for image analysis. J.O.: analysis and plotting of dopaminergic neurons RNAseq. P.G. and M.B.: conceptualization, carry-through and troubleshooting of mice-related experiments. J.J.: mice behavioral experiments. A.M.M.: generation of mice models. T.S.: conceptualization of mice behavior experiments. A.G.: Study design. J.C.S. & R.K.: Conception and study design, supervision of work, administrative and financial support, data interpretation, manuscript revision. C.S.: Conception and study design, supervision of work, performed and analyzed 3D experiments, data interpretation, writing of manuscript, administrative and financial support. All authors approved the final manuscript.

## Acknowledgements

We thank the LCSB sequencing platform (RRID: SCR_021931) and metabolomic platform for the bulk RNAseq and LC-MS and GC-MS experiments, respectively. We also thank Aurélien Ginolhac for developing the snakemake pipeline and François Massart for technical help in performing mouse genotyping. We acknowledge Wolfgang Wurst and Florian Giesert for hosting Anne-Marie Marzesco within the *Helmholtz Zentrum München* as part of the knowledge transfer program, from which the Miro1 p.R285Q mutant mice were generated. We thank Dr. Wagner Zago, Prothena Bioscience, for gifting the 11A5 antibody.

## Methods

### Maintenance of NESC and midbrain organoids generation

Neuroepithelial stem cells (NESC), derived from human iPSC as described previously [73], were used to generate midbrain organoids (MO). 5 lines have been used in the current study: 3 lines from healthy individuals (Ctrl), 1 line from a PD patient carrying a heterozygous point mutation at the *RHOT1* gene (PD-R272Q), and the corresponding isogenic line in which the *RHOT1* mutation has been corrected by CRISPR/Cas9-mediated gene editing (iCtrl) – Table 1. Midbrain organoids were generated following our previously described procedure [19,74]. Briefly, N2B27 medium, composed of 50:50 DMEM-F12 (Thermo Fisher Scientific, cat. no 21331046) and Neurobasal (Thermo Fisher Scientific, cat. no 10888022), 1% penicillin/streptomycin (Thermo Fisher Scientific, cat. no 15140122), 1% GlutaMAX (Thermo Fisher Scientific, cat. no 35050061), 1:100 B27 supplement without vitamin A (Life technologies, cat. no 12587001) and 1:200 N2 supplement (Thermo Fisher Scientific, cat. no 17502001), was used as the base culture media. NESC were cultured in Geltrex-coated (1:90 dilution in media, Thermo Fisher Scientific, cat. no A1413302) 6-well plates in maintenance media (base culture media further supplemented with 150 µM ascorbic acid (AA, Sigma, cat. no A4544), 3 µM CHIR-99021 (Axon Medchem, cat. no CT99021) and 0.75 µM purmorphamine (PMA, Enzo Life Science, cat. no ALX-420-045) until reaching 80% confluency. NESC cell suspension was then obtained using accutase (Sigma, cat. no A6964) and cell density and viability assessed using Trypan Blue (Invitrogen, cat. no T10282) in an automated cell counter (Countess II, Invitrogen). A total of 9000 cells per well, diluted in 150 µl of maintenance media, were plated in 96-well ultra-low attachment plates (faCellitate, cat. no F202003). Plates were then centrifuged at 100g for 3 minutes at room temperature to promote cell contact and facilitate organoid formation. Organoids were cultured in maintenance media for 10 days with media changes every 2 days. At day 8, organoids were embedded in Geltrex droplets and kept in 24-well plates in dynamic conditions (80 rpm). From day 10 to 16, organoids were cultured in maturation media (base media further supplemented with 200 µM ascorbic acid, 500 µM Dibutyryl-cyclic adenosine monophosphate (dbcAMP, STEMCELL Technologies, cat. no 100-0244), 10 ng/ml human brain derived neurothropic factor (hBDNF, Peprotech, cat. no 450-02), 10 ng/ml glial derived neorotropic factor (hGDNF, Peprotech, cat. no 450-10), 1 ng/ml transforming growth factor-beta 3 (TGF-β3, Peprotech cat. no 100-36E) in the presence of 1 µM purmorphamine, with media changes performed every 2 days. From day 16 onward, organoids were cultured in maturation media with media changes occurring every 3 to 4 days. For assays, such as Seahorse and FACS, due to size limitations, non-embedded organoids were used. Here, organoids were kept in maintenance media for 2 days, then cultured for 6 days in maturation media containing 1 µM purmorphamine (media change every 2 days) and finally kept in maturation media until the assay day, with media changes every 3 to 4 days. To ensure cell quality, all cultures were monthly tested for mycoplasma contamination using the LookOut Mycoplasma PCR Detection Kit (Sigma, cat. no MP0035-1KT).

### Generation of 2D midbrain dopaminergic neurons

iPSC-derived NESC (Ctrl1, PD-R272Q and iCtrl; Table 1) were seeded into Geltrex-coated 6-well plates at the density of 3×10^6^ cells/well, unless stated otherwise. Cells were kept in base culture media supplemented with 1µM purmorphamine (Sigma-Aldrich, cat. no SML0868-25mg), 200µM ascorbic acid, and 100 ng/ml fibroblast growth factor 8b (FGF8b, Peprotech, cat. no 10-25) from day 0 (seeding) up to day 8. Between day 8 and day 10, FGF8b was removed and differentiation media contained purmorphamine (0.5 µM) and ascorbic acid (200 µM) only. Finally, from day 10 onwards, differentiating neurons were kept in maturation media, containing 200 µM ascorbic acid, 500 µM dbcAMP (Santa Cruz, cat. no sc-201567C), 10 ng/ml hBDNF, 10 ng/ml hGDNF, and 1 ng/ml TGF-β3.

### Single cell RNA sequencing (scRNAseq) in midbrain organoids

#### 1. Midbrain organoids dissociation and single-cell isolation

30-days old midbrain organoids from one healthy control (Ctrl2), the PD patient carrying the Miro1 p.R272Q mutation (PD-R272Q) and the respective gene-corrected line (iCtrl) were used for scRNAseq analysis (GEO: GSE237133). A total of 30 embedded organoids per condition were pulled for the analysis. Midbrain organoids were collected from their culture medium and washed with 1x PBS (phosphate-buffered saline, Gibco, cat. no 10010-23). Organoids were digested in 1 ml sCelLive™ Tissue Dissociation Solution (Singleron Biotechnologies, cat. no 1190062) in a 15 ml conical tube (Sarstedt, cat. no 62.5544.003) and placed in a thermal shaker at 50 rpm at 37°C for 30 minutes. The state of dissociation was checked at regular intervals under a light microscope. Following digestion, the suspension was filtered using a 40 µm sterile strainer (Greiner, cat. no 542040). Cells were then centrifuged at 350g for 5 minutes at 4 °C, and the pellets resuspended in 500 µl PBS. Cells were stained with Acridine Orange/Propidium Iodide Stain (Logos Biosystems, cat. no F23001), and the cell number and viability were calculated using LUNA-FX7™ Automated Cell Counter (Logos Biosystems).

#### 2. scRNAseq library Preparation

The scRNAseq libraries were constructed using GEXSCOPE™ Single Cell RNAseq Library Kit (Singleron Biotechnologies, cat. no 4161031) according to the manufactureŕs instructions. Briefly, for each library, the concentration of the single-cell suspension was adjusted to 3×10^5^ cells/ml with PBS, and the suspension was loaded onto an SD microfluidic chip to capture 6000 cells. Paramagnetic beads conjugated to oligodT probes carrying a unique molecular identifier (UMI) and a barcode unique to each bead (from the same kit) were loaded, followed by cells lysis. The beads bound to polyadenylated mRNA were extracted from the chip and reverse transcribed into cDNA at 42 °C for 1.5 hours, followed by cDNA amplification by PCR. The cDNA was then fragmented and ligated to indexed Illumina adapters. The fragment size distribution of the final amplified library was obtained using the Agilent Fragment Analyzer.

#### 3. Library sequencing

The library concentration was calculated using the Qubit 4.0 fluorometer and the libraries were pooled in an equimolar fashion. The single cell libraries were sequenced on the Illumina NovaSeq 6000 using a 2×150-bp (base pair) approach to a final depth of 90 GB per library. The reads were demultiplexed according to the multiplexing index sequencing on Illumina’s BaseCloud platform.

#### 4. Transcriptome data pre-processing

Pre-processing of Fastq data were conducted using CeleScope® (v.1.3.0; www.github.com/singleron-RD/CeleScope; Singleron Biotechnologies GmbH, RRID SCR_023553) to generate raw data, using default parameters. Low-quality reads were removed. Sequences were mapped using STAR (https://github.com/alexdobin/STAR), and the human reference GRCh38 and genes were annotated using Ensembl 92. The reads were assigned to genes using featureCount (https://subread.sourceforge.net/; RRID SCR_009803) and the cell calling was performed by fitting a negative bimodal distribution and by determining the threshold between empty wells and cell-associated wells, to generate a count matrix file containing the number of Unique Molecular Identifier (UMI) for each gene within each cell.

#### 5. Transcriptome data processing and plotting

scRNAseq data was analyzed using Seurat R toolkit for single cell genomics version 4.2.0^88^ on R version 4.2.2. Cells with unique feature counts lower than 300 and higher than 7000 were removed as low-quality, empty droplets or probable doublets, respectively. Cells with mitochondrial genes higher than 10 or 15% were also filtered out as additional measure for low quality cells. Datasets were merged, log-normalized and integrated based on the 20 principal component analysis (PCA) components using Seurat integration workflow for better identification of shared cellular populations across the datasets [75]. After integration, merged data were scaled to reduce the variance in gene expression across cells. Seven distinct cell populations were identified applying Louvain algorithm modularity optimization with a resolution of 0.15, based on the top 20 principal components. Visualization was done based on their transcriptomic similarities according to the uniform manifold approximation and projection (UMAP) technique [76]. Cell identities were determined using the online tool GeneAnalytics [77], where marker genes of each cell cluster (identified using the *FindAllMarkers* function of Seurat) were given in the tool for cell type identification choosing *in vitro* parameter for brain cells. The identity of each cluster was further validated by the expression pattern of known cell type specific markers. Differentially expressed genes (DEG) were detected using the *FindMarkers* function of Seurat, comparing the PD-R272Q midbrain (ident.1) against the Control (Ctrl) midbrain (ident.2) or PD-R272Q midbrain (ident.1) against the isogenic control (iCtrl) midbrain (ident.2) in the whole dataset and for each cellular population separately. Significantly differentially expressed genes (p.adjust < 0.05 and logfc threshold = 0.25) were selected for further enrichment analysis using MetaCore (version 2022 Clarivate; RRID SCR_008125), and Rstudio (R version 4.2.2) used to visualize genes fold change from the most relevant enriched pathways.

### Midbrain organoids Western blotting

A total of 3 embedded organoids per cell line per organoid generation (batch) were pulled and snap frozen, followed by protein extraction using RIPA buffer (Abcam, cat. no ab156034) supplemented with 1x Complete^TM^ Protease Inhibitor Cocktail (Roche, cat. no 11697498001) and 1x Phosphatase Inhibitor Cocktail Set V (Merck, cat. no 524629). First, organoids were mechanically dissociated by pipetting in 100 µl of supplemented RIPA buffer followed by a 20 minutes incubation on ice. Lysates were then sonicated at 4 °C for 10 cycles of 30 seconds (sec) on, 30 sec off intervals using the Bioruptor Pico (Diagenode). Lysates were centrifuged at 14,000g for 30 minutes at 4°C, and supernatant was collected. Proteins were then quantified using Pierce™ BCA Protein Assay Kit (Thermo Fisher Scientific, cat. no 23225), according to the manufacturer’s instructions. 15 µg of proteins were diluted in 20 µl of 1x loading buffer (62.5 mM Tris (AppliChem GmbH, cat. no 141940), 1.5% Sodium dodecyl sulfate (SDS; GE Healthcare, cat. no 17-1313-01), 8% glycerol (Alfa Aesar, cat. no A16205), 0.03% bromophenolblue (Carl Roth, cat. no T116.1), 0.5 M DL-Dithiothreitol (DTT; Sigma, cat. no 43816-50ml) and then boiled for 5 min at 95 °C. SDS-PAGE using Bolt™ 4-12% Bis-Tris Plus gels, 1.0 mm (Thermo Fisher Scientific, cat. no NW04127BOX or NW04122BOX or NW04120BOX) in either MOPS (Invitrogen™, cat. no B0001-02, MW > 30) or MES (Invitrogen™, cat. no B000202, MW < 30) buffer was used to separate the proteins by mass. Proteins were then transferred into a PVDF membrane, using the iBlot™ 2 Transfer Stacks (Thermo Fisher Scientific, cat. no IB24001 or IB24002) in the iBlot™ 2 Gel Transfer Device (Thermo Fisher Scientific) for 7 minutes. Membranes were fixed with 0.4% paraformaldehyde (Sigma, cat. no 1.00496.5000) at room temperature for 30 minutes, followed by two 10 minutes washes with PBS and 1 hour incubation with 5% skimmed milk (Carl Roth, cat. no T145.4). Membranes were incubated overnight at 4°C with primary antibodies (table 2) in 5% BSA (Carl Roth, cat. no 8076.3) in PBS with 0.2% tween (Applichem GmbH, cat. no A4974,0500). The following day, membranes were washed 3x for 15 minutes with PBS-T (PBS with 0.02% Tween) and incubated for 2 hours with either anti-rabbit HRP-linked (VWR, cat. no NA934) or anti-mouse HRP-linked (VWR, cat. no NA931) antibodies at 1:1000 dilution (table 2). Finally, membranes were again rinsed 3x for 15 min with PBS-T and revealed using STELLA 8300 imaging system (Raytest) after incubation with chemiluminescent substrate (Life Technologies, cat. no 34580). Proteins of interest were quantified using Image J software (Wayne Rasband; RRID SCR_003070) and normalized for the housekeeping gene beta-actin.

**Table 2.** Antibodies list used for Western blotting and immunostaining experiment in midbrain organoids (MO), dopaminergic neurons (DAN) and mice. aa: amino acid.

| Antibody | Species | Company | Cat. no | RRID | WB Dilution | IF Dilution | Model |
| --- | --- | --- | --- | --- | --- | --- | --- |
| **β-Actin** | Mouse | Cell Signaling | 3700S | AB_2242334 | 1:50 000<br>1:20 000 | - | MO<br>DAN |
| **DAT** | Rat | Millipore | Ab369 | AB_2190413 | - | 1:1000 | Mice |
| **MAP2** | Chicken | Abcam | ab92434 | AB_2138147 | - | 1:1500 | MO |
| **TH** | Rabbit | Santa Cruz | sc-14007 | AB_671397 | 1:1000 | - | MO |
| **TH** | Rabbit | Abcam | ab112 | AB_297840 | - | 1:1000 | MO |
| **TH** | Rabbit | Millipore | Ab152 | AB_390204 | - | 1:1000 | Mice |
| **TH** | Chicken | Abcam | Ab76442 | AB_1524535 | - | 1:1000 | Mice |
| **TUJ1** | Mouse | BioLegend | 801201 | AB_2313773 | 1: 50 000 | 1:1000 | MO |
| **VDAC** | Rabbit | Cell Signaling | 4661 | AB_10557420 | 1:1000 | - | MO |
| **α-Synuclein (2A7; aa 61-95 )** | Rabbit | Novus Biologicals | NBP1-05194 | AB_1555287 | - | 1:1000 | MO |
| **α-Synuclein (C42; aa 15-** | Mouse | BD Transduction | 610787 | AB_398107 | 1:1000 | - | DAN<br>Mice |
| **123)** |  |  |  |  |  |  |  |
| **α-Synuclein**<br>**P-S129**<br>**(11A5)** | Mouse | Prothena | 1347 | Non<br>commercial | 1:1000 |  | Mice |
| **α-Synuclein**<br>**P-S129** | Rabbit | Abcam | ab51253 | AB_869973 | - | 1:1000 | Mice |
| **anti-Rabbit**<br>**IgG (H+L)**<br>**Alexa**<br>**Fluor™ 488** | Goat | Thermo<br>Fisher<br>Scientific | A-11034 | AB_2576217 | - | 1:1000 | MO<br>Mice |
| **Anti-**<br>**Chicken IgG**<br>**(H+L) Alexa**<br>**Fluor™ 488** | Goat | Molecular<br>probes | A-11039 | AB_142924 | - | 1:1000 | Mice |
| **Anti-**<br>**Chicken IgY**<br>**(H+L) Alexa**<br>**Fluor™ 647** | Goat | Thermo<br>Fisher<br>Scientific | A-21449 | AB_10374876 | - | 1:1000 | MO |
| **Anti-rat IgG**<br>**(H+L) Alexa**<br>**Fluor™ 647** | Goat | Molecular<br>Probes | A21247 | AB_141778 | - | 1:1000 | Mice |
| **anti-Mouse**<br>**IgM Alexa**<br>**Fluor™ 568** | Goat | Thermo<br>Fisher<br>Scientific | A-21043 | AB_2535712 | - | 1:1000 | MO |
| **Anti-rabbit**<br>**HRP** | Donkey | Cytiva | NA934 | AB_772206 | 1:1000 | - | MO |
| **Anti-mouse**<br>**HRP** | Sheep | Cytiva | NA931 | AB_772210 | 1:1000 | - | MO |
| **Anti-mouse**<br>**HRP** | Goat | Novex | A24524 | AB_2535993 | 1:10000 | - | DAN |

### Midbrain organoid immunofluorescence staining

Organoids were fixed overnight in 4% paraformaldehyde at 4 °C under shaking conditions. The following day, midbrain organoids were washed 3x for 15 minutes and embedded in 3% low melting agarose (Biozym Scientific GmbH, cat. no 840100). Seventy micrometer sections were then cut using the Leica VT1000s vibratome and kept at 4 °C in PBS with 0.1% sodium azide until further use. For staining, sections were first blocked and permeabilized in a 5% normal goat serum (Thermo Fisher Scientific, cat. no 10000C) and 0.5% Triton X-100 (Carl Roth, cat. no 3051.3) in PBS with 0.1% sodium azide for 2 hours at room temperature under a shaker. Primary antibodies (table 2), diluted in antibody solution (0.1% sodium azide in PBS, 0.01% triton X-100 and 5% goat serum), were incubated at 4 °C for 48 hours on an orbital shaker. Afterwards, sections were rinsed 3x for 10 minutes with PBS containing 0.01% triton X-100 followed by a 2 hours incubation at room temperature under shaking conditions with the respective secondary antibodies (table 2) and the nuclear marker Hoechst-33342 (Invitrogen, cat. no 62249) diluted 1: 10,000 in antibody solution. Finally, organoids sections were further rinsed 3x for 10 minutes with 0.01% triton X-100 PBS and once with miliQ water and, mounted with Fluoromount-G® mounting media (Southern Biotech, cat.no 0100-01) on DBM Teflon® Slides (De Beer Medicals, cat.no BM-9244).

To assess apoptosis, TUNEL assay was performed using the *In Situ* Cell Death Detection Kit, TMR red (Merck, cat. no 12156792910) in combination TH staining. For this, the abovementioned protocol was followed except the secondary antibody incubation. Secondary antibodies (table 2) and Hoechst-33342 were diluted in the TUNEL kit components (label solution with enzyme solution, 1:9) and incubated for 1 hour at room temperature under an orbital shaker.

### Midbrain organoid image acquisition and analysis

Organoids were imaged and analyzed following a pipeline previously described by us [31,32]. Briefly, Yokogawa CV8000 standalone high content screening confocal microscope was used. Cell Voyager and Wako software allowed the automated imaging of organoids section in the x, y and z fields with a 20x objective. Images were then analyzed using adapted in-housed developed MATLAB scripts (v.2021a; RRID SCR_001622). For all experiments, at least one section from 2 organoids per cell line from 3 independent organoid derivations (batches) were quantify unless stated otherwise.

### Midbrain organoids Mito stress test (Seahorse XF)

Seahorse XF Cell Mito Stress Test (Agilent) allowed the evaluation of organoids metabolic state using the Seahorse XFe96 Spheroid FluxPak (Agilent, cat. no 102905-100) in 35 day-old non-embedded organoids. The assay measures oxygen consumption rate (OCR) over a period of time under a specific order of mitochondrial respiration modulators: oligomycin (complex V inhibitor, which inhibits the electron transport reducing OCR), FCCP (carbonyl cyanide-p-trifluoromethoxyphenylhydrazone, uncoupling agent that disrupts proton gradient, when administrated after oligomycin leads to maximal OCR consumption), and finally, rotenone and antimycin (complex I and III inhibitors, respectively, that block any mitochondrial respiration). First, the cartridge containing the O_2_ and pH sensors was hydrated with 200 µl of calibrant solution and incubated overnight at 37 °C in a non-CO_2_ incubator. On the assay day, spheroids microplates were coated with Corning® Cell-Tak™ Cell and Tissue Adhesive (1:60 dilution in 0.1 M sodium bicarbonate; Corning, cat. no 354240) for 1 hour at 37 °C, followed by 2 washes with warm miliQ water. Organoids were then seeded on the spheroid microplates and incubated for at least 1 hour in a non-CO_2_ incubator in Seahorse XF DMEM medium, pH 7.4 (Agilent, cat. no 103575-100) further supplemented with 1 mM L-glutamine (Gibco, cat. no 25030-024), 1 mM pyruvate (Thermo Fisher Scientific, cat. no 11360070), and 21.25 mM glucose (Sigma, cat. no G7021-100g). Cartridge was loaded with assay drugs: 5 µM oligomycin (Sigma, cat. no 75351-5mg), 1 µM FCCP (Abcam, cat. no ab120081), 1 µM antimycin A (Abcam, cat. no ab141904) and 1 µM Rotenone (Sigma, cat. no R8875-1G); and the assay run in Seahorse XFe96 analyzer. Data analysis was done using Seahorse Wave Desktop software (Agilent, RRID SCR_014526). OCR results were normalized by the respective area of each organoids, obtained using Cytation5M reader (BioTek). Datapoints which did not respond to drugs or presented abnormal OCR values were excluded from the analysis. A total of at least 3 organoids per cell line from at least 3 independent organoid generations were used.

### Midbrain organoid flow cytometry

#### 1. Mitochondrial ROS (MitoSox Red)

5 to 10 non-embedded 30 day-old organoids, per cell line, were pulled into 1 well of a 24-well plate and dissociated into single cells with accutase. Replicates were obtained from pulling different independently culture organoids. For organoid dissociation, pulled organoids were incubated with 300 µl of accutase for 1 hour at 37 °C under dynamic conditions, followed by mechanical dissociation with 1000 µl pipette and 200 µl pipette. Incubation with accutase was repeated for 15 to 25 minutes followed by further trituration with a 200 µl pipette. Cell suspension was then collected, centrifuged at 400g for 5 minutes and wash once with assay media. Samples were then stained with 1 nM MitoSOX™ Mitochondrial Superoxide Indicators (Invitrogen™, cat. no M36008) in phenol-red free DMEM-F12 in combination with live-dead stain Zombie NIR (1: 10,000 dilution; Biolegend, cat. no 423106) and incubated for 25 minutes at 37 °C and 5% CO_2_ conditions. Flow cytometer BD LSRFortessa (BD Biosciences) using the BD FACSDiva™ Software (BD Biosciences, RRID SCR_001456) was used to record 10,000 single-cell events per sample. Events counts and intensity were analyzed with FlowJo software (v.10.8.1; RRID SCR_008520). Organoids from at least 3 independent derivations were analyzed.

#### 2. Mitochondrial membrane potential (TMRM & Mitotracker Green)

Organoid single cell suspension was obtained as described above. Once cells were washed in assay media followed by incubation for 30 minutes at 37 °C and 5% CO_2_ with base media containing 1 nM tetramethylrhodamine, TMRM (Invitrogen, cat. no I34361), 100 nM MitoTracker Green (Invitrogen, cat. no M7514) and 1: 10,000 Zombie NIR. Cells were then 2x rinsed in PBS by 400g centrifugation for 5 minutes. Cells were resuspended in PBS and 10,000 single-cell events per sample were recorded at BD LSRFortessa using the BD FACSDiva™ Software. The number and intensity of the events that are double positive for TMRM and Mitotracker Green within the live cell population were analyzed by using FlowJo software (v.10.8.1). Values were obtained from at least 3 independent organoid derivations in 30 days old midbrain organoids.

### Midbrain organoids metabolomics

Polar intracellular metabolites were analyzed in 30 day-old organoids. 5 embedded organoids per line were pulled into Precellys tubes, washed 3 times in miliQ sterile water, snap frozen and kept at −80 °C until processed. A total of 5 independent samples coming from 5 pulled organoids from 2 independent organoid derivations (2 batches) were analyzed. Metabolite extraction and relative quantification were done in a blind way.

#### 1. Metabolite extractions

Metabolite extraction of batch 1 and batch 2 were done separately. Snap frozen organoids were transferred to 2 ml Precellys tubes and 1600 µl (batch 1) or 1500 µl (batch 2) of cold extraction fluid was used. Precellys tubes were prefilled with 600 mg of ceramic beads (1.4 mm, Qiagen, cat. no 1103955) to facilitate metabolite extraction. Pre-cooled (4°C) extraction fluid was composed of 4:1 ratio of methanol (Carl Roth, cat. no AE71.1; Rotisolv, purity ≥ 99.95%) and Mill-Q water (in-house, Milli-Q Advantage A10, 18.2 MΩ•cm, <3 ppb TOC) with 0.8 µg/ml internal standards: internal standards, [_13_C^10^,_15_N^5^] AMP sodium (Sigma-Aldrich, cat. no 650676), 6-chloropurine riboside (Sigma-Aldrich, cat. no 852481), 2-Chloroquinoline-3-carboxylic acid (Sigma-Aldrich, cat. no 688517), 4-Chloro-DL-phenylalanine salt (Sigma-Aldrich, cat. no C6506), Nε-Trifluoroacetyl-L-lysine (Sigma-Aldrich, cat. no 53604), sucralose (Sigma-Aldrich, cat. no 69293) and ^13^C_3_-caffeine-trimethyl (Eurisotop, cat. no CLM-514-1).

Precellys tubes were homogenized with a 30 sec cycle at 6000 rpm (0 to 5 °C). For batch 1, the total volume was divided into 2 Precellys tubes follow by addition of 800 µl of extraction fluid in each followed by a second round of homogenization. A total of 1800 µl of solution from both tubes (batch 1) were transferred to a 2 ml Eppendorf and vortexed (Eppendorf Thermomixer C) for 15 minutes at 2000 rpm (4 °C). For batch 2, after homogenization, 1400 µl of solution was transfer to a 1.5 ml Eppendorf. In both cases, Eppendorf tubes were centrifuged at 21,000g for 10 minutes at 4 °C. After, either two times 500 µl (batch 1) or 800 µl (batch 2) supernatant were transfer to new 1.5 mL Eppendorf tubes, which were evaporated overnight at −4 °C on a refrigerated centrifugal vacuum concentrator (CentriVap 7310000, Labconco). The dried metabolite pellets were stored at −80 °C until LC-MS analysis.

#### 2. Hydrophilic Interaction Liquid Chromatography-Mass Spectrometry (HILIC-MS) measurements

Targeted HILIC-MS measurement was done using a Themo UHPLC Vanquish UHPLC equipped with a binary pump and coupled to a Thermo Exploris 240 mass-spectrometer. Dried metabolite pellets were reconstituted in 50% ACN and filtered using PHENEX-RC 4 mm syringe filter (Phenomenex, cat. no AF0-3203-52). Metabolites were separated using a Hydrophilic Interaction Liquid Chromatography (HILIC) column (SeQuant ZIC pHILIC, 5 μm particles, 2.1 x 150 mm) protected with a guard column (SeQuant ZIC-pHILIC Guard 20 x 2.1 mm). Detailed LC-MS setting are provided in Supplementary Material 1. Peak areas were integrated and exported to Microsoft Excel via the Thermo TraceFinder software (version 5.1; RRID SCR_023045). Metabolite identification confidence was level 1^92^ with all metabolites being verified by an in-house library of standards. Raw values were exported using the sum of all peak areas within a sample. Normalized data was further analyzed using RStudio software (Version 4.3.0; RRID SCR_000432) and further normalized between the different cell lines, to account for cell number differences between conditions, using the Perform Probabilistic Quotient Normalization (PQN).

### Midbrain organoids LDH assay

Cell death was evaluated in 30 day-old non-embedded organoids using LDH-Glo™ Cytotoxicity Assay (Promega, cat. no J2380) accordingly to the manufacturer’ instructions. Briefly, organoid media was changed to freshly prepared media 24 h before the assay. For the assay, 25 µl of media from at least 2 organoids per cell line from 3 independent organoid derivation were used. 25 µl of LDH detection reagent (25 µl LDH Detection Enzyme Mix with 0.125 µl Reductase Substrate) were added to the 25 µl of organoid media and incubated in lumitrac-600 96-well plate (Greiner, cat. no 655074), at room temperature, for 1 hour under shaking at 400 rpm using a ThermoMixer C (Eppendorf). Luminescent was read at the Cytation5M reader (BioTek). Relative levels of LDH were calculated by LDH luminescence normalized by organoid area set to 100.

### Bulk RNA sequencing in dopaminergic neurons

RNA extraction was performed using the RNeasy Kit (Qiagen, cat. no 74104) according to manufacturer’s instructions in dopaminergic neurons after 30 days of differentiation using 600 µl RLT lysis buffer containing 1:1,000 2-mercaptoethanol (Sigma Aldrich, cat. no M3148). After RNA extraction, eluted RNA concentration was assessed using a NanoDrop™ spectrophotometer. RNA quality was further assessed using the Agilent 2100 Bioanalyzer showing RNA integrity (RIN) values > 8.

RNA library preparation was performed using TruSeq Stranded mRNA library prep kit (Illumina, cat. no 20020594) according to the manufacturer’s protocol, and then sequenced using NextSeq2000 (Illumina) at the LCSB Genomics platform (RRID SCR_021931). For all samples paired end reads of 51 bp length were generated.

Data (GEO: GSE238129) was processed using an in-house snakemake workflow (https://git-r3lab.uni.lu/aurelien.ginolhac/snakemake-rna-seq; release v0.2.3 and singularity image v0.4). Snakemake pipeline is under license https://git-r3lab.uni.lu/aurelien.ginolhac/snakemake-rna-seq/-/blob/main/LICENSE. Raw read quality was assessed by FastQC (v0.11.9; RRID SCR_014583) [78]. Adapters were removed using AdapterRemoval (v2.3.2; RRID:SCR_011834) [79], with a minimum length of the remaining reads set to 35 bp. Reads were mapped to hg38 (GRCh38.p13) using STAR (v.2.7.9a) [80], and reads were counted using featureCounts from the R package Rsubread (v2.8.1; RRID SCR_016945) [81]. All transcripts with counts > 10 were used for differential gene expression analysis using the R package DESeq2 (v1.34.0; RRID SCR_015687) [82]. apeglm was used for normalization in DEseq2 (v.1.16.0) [83]. FPKM (fragments per kilobase of exon per million mapped fragments) were calculated using DESeq2 package. Pathway analysis on DEG with minimum FPKM of > 1, false discovery rate < 0.05, and a minimum log2-fold change cut-off of +/−1.5 was performed using Ingenuity Pathway Analysis tool (Qiagen, version: 60467501) and the EnrichR online tool (https://maayanlab.cloud/Enrichr; RRID SCR_001575).

### Dopaminergic neurons imaging

iPSC-derived dopaminergic neurons were re-plated on day 15 of differentiation at a density of 100,000 cells per well into PerkinElmer Phenoplate 96-well plates (PerkinElmer, cat. no 6055300) and kept in maturation media until imaged (day of differentiation 30, approximately). Yokogawa CV8000 standalone high content screening microscope using a 60x (intracellular ROS and calcium imaging) or 20x (mitochondrial membrane potential) objective was used for dopaminergic neurons live-cell imaging experiments. To maintain cell integrity during image acquisition, neurons were kept under controlled conditions: 5 % CO_2_, 37 °C temperature and 80% humidity. A total of 17 fields for intracellular ROS, 1 field for calcium imaging, and 21 fields in the x, y, z axis were acquired per well. MATLAB scripts (v.2021a) developed in-house were further used for image analysis.

#### 1. Intracellular ROS (CellROX Deep Red & CellTracker Green)

The cell-permeant CellROX Deep Red dye (Thermo Fisher Scientific, cat. no C10422), which allows for live assessment of intracellular ROS, was used in combination with the general cellular marker CellTracker Green (Thermo Fisher Scientific, cat. no C7025), and nuclei dye Hoechst-33342 (Invitrogen, cat. no 21492) to evaluate intracellular ROS using live imaging. Two days before the assay cells were changed to maturation media without antioxidants (without B27 and ascorbic acid). At the day of the assay, dopaminergic neurons were incubated for 30 minutes with 10 µM CellROX Deep Red, 0.5 µM CellTracker Green and 1 µg/ml Hoechst-33342 at 37 °C with 5% CO_2_, washed once with PBS and kept in maturation media during imaging acquisition.

#### 2. Mitochondrial membrane potential (TMRE & Mitotracker Green)

Dopaminergic neurons were stained with the specific mitochondria membrane potential maker TMRE (Invitrogen™, cat. no T669) at concentration of 20 μM, combined with 0.1 nM MitoTracker Green (Invitrogen™, cat. no M7514) and 1 µg/ml Hoechst-33342 in base culture media for 30 minutes in a cell culture incubator (37 °C with 5% CO_2_). Cells were washed once in PBS and further re-incubated in base culture media with 20 μM TMRE followed by live imaging.

#### 3. Calcium imaging

For calcium imaging, dopaminergic neurons were incubated for 1 hour with 50% maturation media 50% of 2X calcium indicator Fluo4-Direct (Thermo Fisher Scientific, cat. no F10471) following the manufacturer’s instructions, and Hoechst-33342 (1 µg/ml). Imaging was done for a total of 10 minutes at 0.5 Hz. For the first minute, imaging was done at basal conditions to establish baseline calcium levels (F0). At minute 1, dopaminergic neurons were exposed to the calcium ionophore ionomycin (10 μM; Sigma-Aldrich, cat. no I0634-1MG), dispensed in an automated way by Yokogawa CV8000, allowing to understand dopaminergic neurons response to calcium influx increase (F1).

### Dopaminergic neurons Seahorse mitochondrial stress test

OCR and pH was measured in whole cells using the Seahorse Xfe96 Cell Metabolism Analyzer (Agilent) and Seahorse FluxPak (Agilent, cat. no 103775-100) following Agilent Seahorse mito stress test. Dopaminergic neurons were re-plated at a density of 100,000 cells per well, in maturation media, into Geltex-coated Seahorse Xfe96 well plate 24 hour prior to the assay. After seeding, the plate was left at room temperature for 1 hour to prevent edge-effects. Outer wells were filled with media only and not used for cells seeding to avoid cell stress occurring due to evaporation. Dopaminergic neurons were then kept in a cell culture incubator overnight. Also, at day before the assay, XFe96 sensor cartridge was hydrated using 200 μl of XF-calibrant solution. Seahorse base medium (Agilent, cat. no 102353) was further supplemented with 382 µg/ml D-glucose (Sigma, cat. no D8375), 2 mM (1%) L-glutamine (Gibco, cat. no 35050061) and 40 µg/ml sodium pyruvate (Sigma, cat. no P5280; pH = 7.4 ± 0.05 at 37 °C). Both, cartridge and media were incubated overnight at 37 °C in a non-CO_2_ incubator.

At the assay day (day of differentiation 30), the media of dopaminergic neurons was replaced by Seahorse assay media by removing 80 μl of maturation media (leaving 20 µl) from each well, rinsed 2x with 200 μl of Seahorse assay media and finally adding 155 μl of Seahorse assay media per well to a total final volume of 175 μl/well. The plate was then incubated in a non-CO_2_ incubator for at least 1 hour to allow cells to equilibrate. Mitochondria targeted drugs dissolved in Seahorse assay media (25 µl) were loaded into the cartridge: 1 µM oligomycin, 1 µM FCCP (Sigma, cat. no C2920), 0.5 µM antimycin A (Sigma, cat. no A8674) and 0.5 µM Rotenone. After, Mito stress test assay was run in the Seahorse Xfe96 Cell Metabolism Analyzer and OCR and pH measure through time.

Assay normalization was done by DNA quantification using the CyQUANT® assay (Invitrogen, cat. no C7026). For that, the Seahorse assay media was removed and dopaminergic neurons were frozen and stored at −80 °C in the Seahorse cell culture plate until DNA quantification. CyQUANT® components as well as dopaminergic neurons were brought to room temperature. The CyQUANT® GR stock solution was diluted 400-fold into the 1X cell-lysis buffer, 200 μl of solution were added to each well and incubated for 2–5 minutes at room temperature, protected from light. Samples were then transferred into a new 96-well plate and fluorescence measure by Cytation5M reader at 480 nm excitation and ∼520 nm emission.

### Dopaminergic neurons NAD(P)/NAD(P)H and ATP measurement

At day of differentiation 30, dopaminergic neurons were harvest with accutase for ATP measurement (40,000 cells per replicate), NAD(P)/NAD(P)H (25,000 cells per replicate). Measurements were done using commercially available kits from Promega, following the manufacturer’s protocol: CellTiter-Glo (Promega, cat. no G7570), NAD/NADH-Glo (Promega, cat. no G9071) and NADP/NADPH-Glo (Promega, cat. no G9081). Luminescence was read using the Cytation5M reader. All values were normalized to protein amount quantified with the Pierce™ BCA Protein Assay Kit according to manufacturer instructions. For all assays, a total of 4 independent batches (derivations) with 2 technical replicates per batch were performed.

### Extracellular metabolomics on dopaminergic neurons

#### 1. Sample collection and metabolite extraction

At day 30 of differentiation, maturation media that has been in contact with dopaminergic neurons for the previous 48 hours was collected, filtered using Phenex Regenerated Cellulose (RC) syringe filters to remove any cells or debris, and frozen at −80 °C until extraction and gas chromatography (GC)-MS measurement. Metabolite derivatization was performed by using a multi-purpose sample preparation robot (Gerstel). Dried medium extracts were dissolved in 30 µl pyridine, containing 20 mg/ml methoxyamine hydrochloride (Sigma-Aldrich, cat. no 89803), for 120 min at 45 °C under shaking. After adding 30 µl of N-methyl-N-trimethylsilyl-trifluoroacetamide (Macherey-Nagel, cat. no 701270.510) samples were further incubated for 30 minutes at 45 °C under continuous shaking followed by GC-MS analysis.

#### 2. Gas chromatography – Mass spectrometry (GC-MS)

GC-MS analysis was performed on an Agilent 8890 GC coupled to an Agilent 5977B MS (Agilent Technologies). A sample volume of 1 µl was injected into a Split/Splitless inlet, operating in split mode (20:1) at 270 °C. The gas chromatograph was equipped with a 30 m (I.D. 0.25 mm, film 0.25 µm) ZB-5MSplus capillary column (Phenomenex, cat. no 7HG-G030-11-GGA) with 5 m guard column in front of the analytical column. Helium was used as carrier gas with a constant flow rate of 1.4 ml/minute. The GC oven temperature was held at 90 °C for 1 minute and increased to 220 °C at 10 °C/minute. Then, the temperature was increased to 300 °C at 20 °C/minute followed by 4 min post run time at 325 °C. The total run time was 22 minutes. The transfer line temperature was set to 280 °C. The MSD was operating under electron ionization at 70 eV. The MS source was held at 230 °C and the quadrupole at 150 °C. Mass spectra were acquired in selected ion monitoring (SIM) mode for precise quantification of medium components. Supplementary Material 2 shows the masses used for quantification and qualification of the derivatized target analytes (dwell times between 20 and 70 ms).

#### 3. Data processing and normalization

All GC-MS chromatograms were processed using MetaboliteDetector (v3.2.20190704) [84]. Compounds were annotated by retention time and mass spectrum using an in-house mass spectral (SIM) library (overall similarity > 0.80). The following deconvolution settings were applied: peak threshold: 2; minimum peak height: 2; bins per scan: 10; deconvolution width: 8 scans; no baseline adjustment; minimum 1 peak per spectrum; no minimum required base peak intensity. The internal standards (U-13C5-ribitol and pentanedioic-d6 acid) were added at the same concentration to every sample to correct for uncontrolled sample losses, and analyte degradation during metabolite extraction and sensitivity drifts during measurements. The dataset was normalized by using the response ratio of the integrated peak area of the analyte and the integrated peak area of the internal standard. Further normalization based on the total cell number that the media was in contact with was performed and the relative abundance of metabolites was plotted in GraphPad Prism v10 (RRID SCR_002798) using negative values for metabolites consumed and positive values for metabolites released by the dopaminergic neurons.

### Dopaminergic neurons Western blotting

Neurons were lysed with 200 μl SDS lysis buffer (1% SDS with protease inhibitor cocktail tablet; Roche, cat. no 04693159001) per well in 6-well plates and scraping. Lysates were transferred into 1.5 ml tubes and boiled for 5 minutes at 95 °C. Protein quantification was performed using the Pierce BCA assay kit according to manufacturer’s instructions. NuPAGE™ 4 to 12%, Bis-Tris, 1.0 mm, Mini Protein Gels with 12-wells were used for all blots (Invitrogen™, cat. no NP0329PK2). Prior to loading, samples were diluted with 6x Lämmli loading buffer and boiled for 5 minutes at 95 °C. As ladder, PageRuler Plus Prestained Protein Ladder was used (Thermo Fisher Scientific, cat. no 26619). Electrophoresis were runed in NuPAGE™ MES SDS running buffer. After the run, wet transfer ran for 75 minutes at 100 V in transfer buffer: 1:5 99% ethanol and 1x Tris glycine in MilliQ water. After protein transfer, Ponceau red was used to assess total protein levels. The membrane was then rinsed with TBS-T (Tris-buffered saline with 0.1% Tween 20 detergent) and blocked with 5% BSA in TBS-T for 1 hour prior to incubation of the primary antibody overnight at 4 °C with rotation: α-synuclein (C42) and β-actin (8H10D10) – Table 2. The next day, membrane was washed 3x for 10 minutes in TBS-T with shaking and incubated with the respective secondary antibody (Table 2) for 1 hour at room temperature with shaking. Then, membrane was further washed 3x for 10 minutes in TBS-T with shaking. Protein bands were detected with ECL Prime Western blotting Amersham (Sigma Aldrich, cat. no GERPN2232) detection reagent and images acquired with STELLA imaging system. Western Blot membranes were analyzed with Image J software.

### In vivo

All mouse experiments were performed according the European FELASA guidelines for animal experimentation (see ethical approval section).

#### 1. Housing

Mice were housed and bred in dedicated facilities under specific pathogen free (SPF) conditions, then transferred to a conventional mouse facility at least one week before any manipulation or measurement. Both facilities had 12 hours alternating dark/light cycles. Mice always had *ad libitum* access to standard mouse food (Sniff, cat. no V 1534-300) and water. Health monitoring was performed quarterly and yearly, according to the FELASA 2014 pathogen list.

#### 1. Generation of Miro1 p.R285Q knock-in mice

Miro1 p.R285Q knock-in mice (which represent the human Miro1 p.R272Q ortholog mutation) were generated by CRISPR/Cas9-mediated gene editing in mouse zygotes. Briefly, pronuclear stage zygotes were harvested from C57BL/6N mice super ovulated C57BL/6N females mated with C57BL/6N males. The pronucleus of embryos were microinjected with a mix containing 25 ng/µl Cas9 mRNA, 60 ng/µl Recombinant S. pyogenes Cas9 nuclease, 0.6 pmol/µl Alt-R® CRISPR-Cas9 crRNA (protospacer TGGACTGTGCTTCGACGATT; IDT, Inc.): 0.6 pmol/µl Alt-R® CRISPR-Cas9 tracrRNA duplex (IDT, Inc.) and 50 ng/µl of specific mutagenic ssODN Miro1 R285Q (5’ - GGTTTTCTCTTTTTACATACACTTTTTATCCAGAGGGGGAGGCATGAGACTACTTGGACTGTGCTTCAGCGGTTTGGTTATGACGATGACCTGGACCTGACGCCTGAGTATTTATTCCCCCTGTATGTACCTCAGCGCTC - 3’, synthetized by Metabion international AG), comprising the R285Q substitution and an additional silent mutation for genotyping purposes. After microinjections, zygotes were transferred into pseudopregnant CD-1 foster mice.

To identify Miro1 p.R285Q knock-in founder mice (F0 generation) derived from microinjected zygotes, genomic (g)DNA was isolated from tissue biopsies from mice at the age of 3 - 4 weeks and used to assess Miro1 p.R285Q substitution by PCR (see *Mice genotyping* section below). The knock-in founder mice (F0) were generated at the Institute of Developmental Genetics (Helmholtz Zentrum München, Germany), according to protocols approved by the government of Upper Bavaria and in accordance with the guidelines of the European Community Council Directives.

The animals of Miro1 p.R285Q knock-in mouse line (B6.Miro1^tmR285QHmgu^) were raised and handled at the University of Luxembourg following the European Union directive 2010/63/EU (see ethical approval section). Wild-type (wt/wt) were used to generate heterozygous (wt/R285Q) and homozygous (R285Q /R285Q) Miro1 p.R285Q mutant mice (B6.Miro1^tmR285QHmgu^), which represent the human Miro1 p.R272Q ortholog mutation, using an heterozygous x heterozygous and homozygous x homozygous breading strategy.

#### 2. Mice genotyping

Genomic DNA from biopsies was extracted using the Nucleospin DNA RapidLyse kit (Macherey Nagel, cat. no 740100.250) and quantified using a NanoDrop™ spectrophotometer. DNA was then used to assess Miro1 p.R285Q point mutation integration by PCR. 300 ng of gDNA and 0.2 µM Miro1 forward and reversed primers (Table 3) were used in a total of 50 µl reaction. PCR reaction was done in Biometra Thermocycler T Professional Basic 96 (Montreal Biotech, cat. no 846-070-701) for 5 minutes at 94 °C, followed by 35 cycles of 1 minute at 94 °C denaturation step, 1 minute at 60 °C annealing and 3 minutes 72 °C elongation step, and a final extension step of 7 minutes at 72 °C. PCR products were further digested by TAQI (GoTaq® G2 Flexi DNA Polymerase; Promega, cat. no M7805) to assess if the point mutation is or not present, since its restriction site will be only present in wt Miro1 protein. For that, 12 µl of ddH2O, 10 µl PCR product (no purification is necessary), 0.5 µl NEB Taq1 enzyme and 2.5 µl 10x smart cut NEB Buffer (Bioke cat. no R0149S) were amplified by PCR using the thermocycling program: 40 minutes at 65 °C digestion, 20 minutes 80 °C inactivation step, and a final hold at 7 °C. PCR products and the GeneRuler^TM^ 100 bp Plus DNA ladder were then ran in 1.5% agarose gel at 130 V for 45 minutes and revealed in a Biodoc Analyse GBOX machine (Syngene) under UV light using Midori Green advance (Biozyl, cat. no 617004). Considering the primers used, wt Miro1 will present 2 bands at 334 bp and 569 bp, while homozygous Miro1 p.R285Q mutant will only present one band with 903 bp.

**Table 3.**
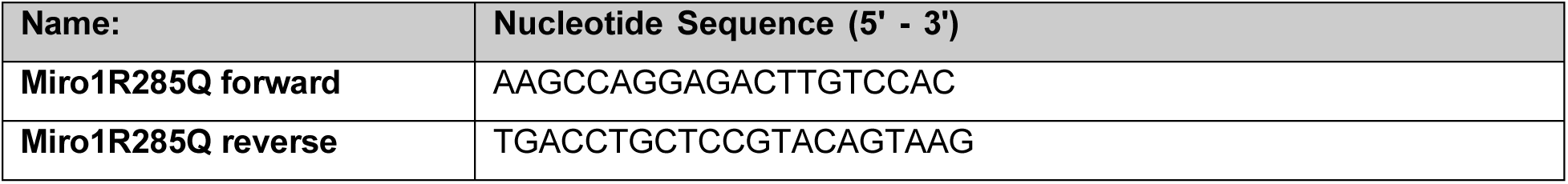
PCR primers used for PCR of mice.

### Mice Behavior

Mice weight was assessed throughout the study at month 12, 15, 18 and 21.

#### 1. Home cage activity

The LabMaster system (TSE Systems) was used to record and analyze the spontaneous home cage activity of the mice. The mouse cage was surrounded by sensor frames with which the number of beam brakes was quantified and analyzed in 15 minutes intervals. The recorded data provided information on total activity, movement and rearing. Additionally, the volume of consumed food and water was quantified. The data was recorded over a period of 22 hours, which started with about 30 minutes of light phase before a 12 hour phase of darkness. The measurement happened in the same day and night cycle that mice were used to. All mice were analyzed in individual cages and afterwards reunited with their cage mates. Measurements were done in 20 month-old mice.

#### 2. RotaRod

An accelerating rotarod (TSE Systems) was used to measure the motor coordination abilities, balance and spontaneous motor activity/movement anterograde procedural memory of the mice. The analysis was performed in a room with only red light illumination during the night time of the animals. Mice were placed on the rod which accelerated from 4 to 40 rpm over a duration of 300 sec. The latency to fall off the rotating rod was recorded in each trial. If a mouse fell within the first 10 sec of a trial the measurement was repeated. Per test day three trials were carried out with a minimum 15 minutes rest between trials. The mean latency to fall per mouse was calculated as the mean value of the three trials. Rotarod was performed twice with the same female mice at age 20 and 21 months. There was no training prior to the first trial. One female was excluded based on their significant higher weight.

### Mice brain dissection

Mice were deeply anesthetized with ketamine-medetomidine anesthesia (150 mg/kg ketamine lot nr. 136472 and 1 mg/kg medetomidine lot nr. 2033.85) and transcardiacally perfused with PBS. After perfusion, brains were placed on ice, and split along the longitudinal fissure into the two hemispheres. For molecular biology analyses, one part was dissected into different regions of interest (striatum and midbrain), put on dry ice and stored at −80 °C. The other hemisphere were fixed for immunohistochemistry in 4% PFA for 48 hours at 4 °C and then stored in PBS with 0.2% sodium-azide until further processing.

### Immunohistochemistry (mice)

Female and male young (3 to 6 month-old) and aged (15 month-old) mice were used for analyzing striatal DAT and TH as well as SNpc TH levels. Serialized parasagittal free floating, 50 µm-thick sections were obtained using a Leica VT-1000S vibratome and collected in cryoprotective medium composed of a 1:1 ethylene glycol (Sigma Aldrich, cat. no 61941) and PBS solution, supplemented with 1% w/v polyvinyl pyrrolidone (Sigma Aldrich, cat. no PVP40). Sections were stored at −20 °C, in tubes, each containing series of every 4^th^ section.

For staining, sections were washed 2x 5 minutes in washing buffer (PBS with 0.1% Triton X-100) with agitation, permeabilized for 30 minutes in PBS containing 1.5% Triton X-100 and 3% hydrogen peroxide, to deactivate endogenous peroxidases. Sections were further washed 2x 5 minutes in washing buffer and blocked in PBS with 5% BSA and 0.1% Triton X-100 for 30 minutes. After, sections were rinsed 2x 5 minutes in washing buffer followed by incubation with primary antibodies: rabbit anti-TH, rat anti-DAT Nter for the striatum; chicken anti-TH for SNpc (Table 2) in antibody buffer (PBS with 0.3% Triton X-100 and 2% BSA) overnight under strong shaking at room temperature. After three more washing steps (10 minutes each), sections were incubated with appropriate secondary antibodies (Table 2) in antibody buffer, in the dark, on shaking conditions for 2 hours at room temperature. Next, the sections were washed again (3x 10 minutes) in washing buffer, and once in TBS (Trizma base 6.06 g/l (Sigma Aldrich, cat. no T1503), 8.77 g/l NaCl (Sigma Aldrich, cat. no S9888), adjusted with HCl to pH 7.4). Finally, sections were mounted on glass slides, cover slipped using Dako fluorescent mounting medium (DAKO, cat. no S302380).

### Imaging acquisition and quantification (mice)

Imaging was done using a Zeiss *AxioImager Z1* upright microscope, coupled to a “*Colibri*” LED system to generate fluorescence light of defined wavelengths, an *Mrm3* digital camera for image capture, and equipped with a PRIOR motorized slide stage. The complete Zeiss imaging system was controlled by Zeiss Blue Vision software. To measure TH and DAT, 2-3 striatal sections per mouse containing 3 fields in the dorsal striatum were taken in a 40X objective using the Apotome. Apotome captured image were then modified with the parameters “display optical sectioning” to correct phase errors. Obtained images were converted to TIFF. Then, after thresholding, the percent area occupied by TH and DAT staining were determined using the publicly available imaging software FIJI (RRID SCR_002285), using the “restrict to threshold” parameter. Qualitative assessment of phosphor S129 α-synuclein was performed on 15 µM deep stacks at a 40 % magnification, followed by maximum intensity projection.

For TH quantification, we first carry a careful anatomical observation to distinguish SNpc specific TH-positive neurons from the ones present in other anatomical regions. We recognized the SNpc at four anatomically distinguishable levels using 6-10 sections of 50 µm (spaced 200 μm) per mouse, covering the entire space occupied by the SNpc in each mouse brain. Then, for each section, tiled pictures (2×2) were taken at 10x magnification using the above-mentioned Zeiss imaging system. Next, the region-of-interest (ROI) tool in the FIJI software was used to segment the area occupied only by SNpc TH-positive neurons within each section. After thresholding, the area occupied (in pixels) by TH-positive neurons was measured. The four distinguishable anatomical levels of the SNpc were measured using 2-3 sections/level in each mouse. Values for each level were averaged separately. Thus, 4 final area values, representative of the areas occupied by TH-positive neurons in each anatomical level of the SN, were obtained for each mouse. To obtain one representative value for each brain hemisphere, these 4 values were summed up (“cumulated SN surface”), and converted to mm^2^ for graphical representation [85]. Detailed information on the method and its correlation with stereological cell counts can be found in Ashrafi et al. [85].

### Striatal dopamine measurement

Striatum, dissected as abovementioned, was used to quantify the abundance of the neurotransmitter dopamine in 15 month-old mice. Tissue was used to extract and measure dopamine by GS-MS based on previously published method [86,87]. Briefly, 500 µl of methanolic extraction fluid (4:1, methanol/water mixture, v/v) were added to 50 mg mouse brain (striatum). The extraction fluid contained an internal standard mix, consisting of U-13C5 ribitol (c = 2 µg/ml; Omicron Biochemicals, cat. no ALD-062) and pentanedioic-d6 acid (c = 2 µg/ml; C/D/N Isotopes Inc., cat. no D-5227). Samples were subsequently homogenized using a Precellys24 homogenizer (Bertin Technologies) using 600 mg ceramic beads (1.4 mm) and one 30 sec cycle at 6,000 rpm at 0 to 5 °C. Then, 250 µl of a 0.1 mol/l hydrochloric acid solution, including dopamine-D4 (c = 15 µmol/l) was added to the homogenate. Polar metabolites were extracted by adding 400 µl of chloroform. After incubation under shaking for 15 minutes at 2,000 rpm at 4 °C (Eppendorf ThermoMix Comfort), samples were centrifuged for 5 minutes at 21,000g at 4 °C. 80 µl of the upper phase containing the polar metabolites were transferred into a GC glass vial with micro insert and evaporated at −4 °C for 4 hours, followed by an adaptation phase to room temperature for 25 minutes (Labconco CentriVap). Samples were submitted to subsequent GC-MS analysis.

### Data analysis and statistics

GraphPad Prism (version 10) was used to plot and statistically analyzed midbrain organoids LDH assay, organoids and dopaminergic neurons Seahorse data, dopaminergic neurons metabolomics, calcium imaging and ATP, NAD(H) and NADP(H) assays, while RStudio software (version R 4.3.0) was used to plot and statistically analyzed dopaminergic neurons and/or midbrain organoids Western blotting, image analysis and metabolomic data. Data are expressed as mean ± standard error of mean (SEM). For midbrain organoids LDH and dopaminergic neurons single cell calcium imaging assays outlier removal was performed with ROUT test (Q=5%) using GraphPad Prism. For TUNEL assay, Rosner Test (from EnvStats package) was used to detect outliers using RStudio. Statistical significance was determined with GraphPad Prism 10 software or with RStudio. First, data was analyzed for normality using Shapiro test. For normal distributed data (Shapiro test not significant) ANOVA with post-hoc Tukey HSD test was used, while in non-normal data (Shapiro test significant) the statistical significance was calculated using the non-parametric Kruskall-walis test. P < 0.05 was considered to represent statistical significance, with * P < 0.05, ** P < 0.01 and *** P < 0.001.

## Ethical approval

All necessary ethical approvals have been obtained in accordance with the Règlement grand-ducal du 11 janvier 2013 relatif à la protection des animaux utilisés à des fins scientifiques” adapted from and in line with the European Directive 2010/63/EU.w (on the use of clinical samples and on the Care and Use of laboratory animals, where applicable). The Luxembourgish National Research Ethics Committee (CNER) provided ethical approval for the following projects: “Disease modelling of Parkinson’s disease using patient-derived fibroblasts and induced pluripotent stem cells” (DiMo-PD, CNER #201411/05) and, *‘In vitro* modeling of Parkinson’s disease (ivPD)’ (ERP 18-082 ivPD).

## Data availability

All the data used in the current study, both raw and processed, as well as all the scripts developed are publicly available at https://doi.org/10.17881/vm4y-sv50. The scripts generated from the current are also available on gitlab: https://gitlab.lcsb.uni.lu/dvb/saraiva_2023. Midbrain organoid scRNAseq and dopaminergic neurons bulk RNAseq data are also available on Gene Expression Omnibus (GEO) repository under the accession code GSE237133 and GSE238129, respectively.

