## Supplementary Figure for "Parkinson’s disease-related Miro1 mutation induces mitochondrial dysfunction and loss of dopaminergic neurons *in vitro* and *in vivo*"

**Supplementary Figures**

**Supplementary figure 1. Midbrain organoid (MO) single cell RNA sequencing quality control and cell cluster markers. A)** Plots representing the gene number (nFeature_RNA), total molecules number (nCount_RNA), mitochondrial gene percentage as well as the correlation between the number of genes and total molecules before (unfiltered; left) and after (filtered; right) applying the quality control thresholding in **(i)** healthy MO (Ctrl2), **(ii)** Miro1 p.R272Q mutant organoids (PD-R272Q), and **(iii)** isogenic control (iCtrl). **B)** Expression levels of cell specific markers for oligodendrocyte progenitors, glia progenitors, neural progenitors, dopaminergic neurons, other neuronal subtypes and mature neurons, respectively, in the different MO clusters identified.

**Supplementary figure 2. Deregulated process and pathways in midbrain organoids (MO). A)** Top 25 most deregulated process (left) and pathways (right) in Miro1 p.R272Q mutant MO (PD-R272Q) when compared with healthy control (Ctrl) or isogenic control (iCtrl) (PD-R272Q vs Ctrl; PD-R272Q vs iCtrl) based on differential express gene analysis of the single cell RNA sequencing data. **B)** Top 25 most deregulated process (left) and pathways (right) in PD-R272Q vs Ctrl and PD-R272Q vs iCtrl in the dopaminergic neuron clusters (dopaminergic neurons 1 and dopaminergic neurons 2).

**Supplementary figure 3. Dopaminergic Neurons bulk RNA sequencing.** Expression levels in fpkm (fragments per kilobase per million mapped fragments) of dopaminergic neuronal markers (top) and glial markers (bottom) expressed by dopaminergic neurons in the bulk RNA sequencing dataset.

**Supplementary figure 4. Midbrain Organoids deregulated genes from single cell RNA sequencing.** Fold change of the significant differentially expressed genes contributing to the deregulation of the *'oxidative stress ROS-induced cellular signaling'* pathway in midbrain organoids comparing Miro1 p.R272Q midbrain organoids (PD-R272Q MO) and healthy (Ctrl) or isogenic (iCtrl) controls (i.e. PD-R272Q vs Ctrl and PD-R272Q vs iCtrl).

**Supplementary figure 5. *RHOT1* expression levels *in vitro*. A)** Ridge plot of midbrain organoids *RHOT1* gene expression in Miro1 p.R272Q mutant (PD-R272Q), healthy (Ctrl) and isogenic (iCtrl) controls conditions based on the single cell RNA sequencing data. **B)** Ctrl, PD-

R272Q and iCtrl expression levels of *RHOT1* in dopaminergic neuronal cultures in the bulk RNA sequencing dataset.

**Supplementary figure 6. Deregulated gene expression of midbrain organoids using single cell RNA sequencing data.** Significant differentially expressed genes contributing to the deregulation of the process “*Apoptosis_Apoptotic mitochondria*” (left) and the pathway “*Apoptosis and survival_NGF/TrkA PI3K−mediated signaling*” found within the dopaminergic neuron clusters (dopaminergic neurons 1 and dopaminergic neurons 2 clusters) of the midbrain organoids. Graph displays gene fold-changes comparing Miro1 p.R272Q midbrain organoids (PD-R272Q MO) and healthy (Ctrl) or isogenic (iCtrl) controls (i.e. PD-R272Q vs Ctrl and PD-R272Q vs iCtrl).

**Supplementary figure 7: Miro1 p.R285Q mutant C57BL/6 mice features. A)** Body weight of Wilde-type (wt/wt, green) and Miro1 p.R285Q mutant heterozygous (wt/R285Q, orange) and homozygous (R285Q/R285Q, red) mice at month 12, 15, 18 and 21 of age. **B-C)** Quantification of tyrosine hydroxylase (TH; **B)** and dopamine transporter (DAT; **C**) terminals in the striatum of 3 to 6 month-old mice. **D)** TH cumulative area in the substantia nigra (SNpc) of 3 to 6 month-old mice. **B-D)** wt/wt: n = 6F; wt/R285Q: n = 6F; R285/R285Q: n = 10F. **E)** DAT quantification in the striatum of 15 month-old mice. **F)** Graphic depicts striatal dopamine quantification (pmol/mg) in wt/wt and R285Q/R285Q 15 month-old mice using gas-chromatography-mass spectrometry (GC-MS). **15 month-old mice:** wt/wt: n = 2M + 5F; wt/R285Q: n = 3M + 5F; R285/R285Q: n = 4M + 6F. **G)** Quantification of SNpc TH positive area in 15 month-old male mice. **H)** Representative image of pS129 α-synuclein and TH immunoreactivity in the SNpc of old male wildtype and Miro1 p.R285Q homozygous mice. Scalebar 20µm. **J, K)** Consumption of water **(J)** and food **(K)** for 22 hours in 20 month-old mice measured with the Phenomaster® Cage. **L)** Total activity, expressed as number of beam brakes every 15 minutes, for 22 hours, in 20 month-old mice measured with the Phenomaster® Cage.

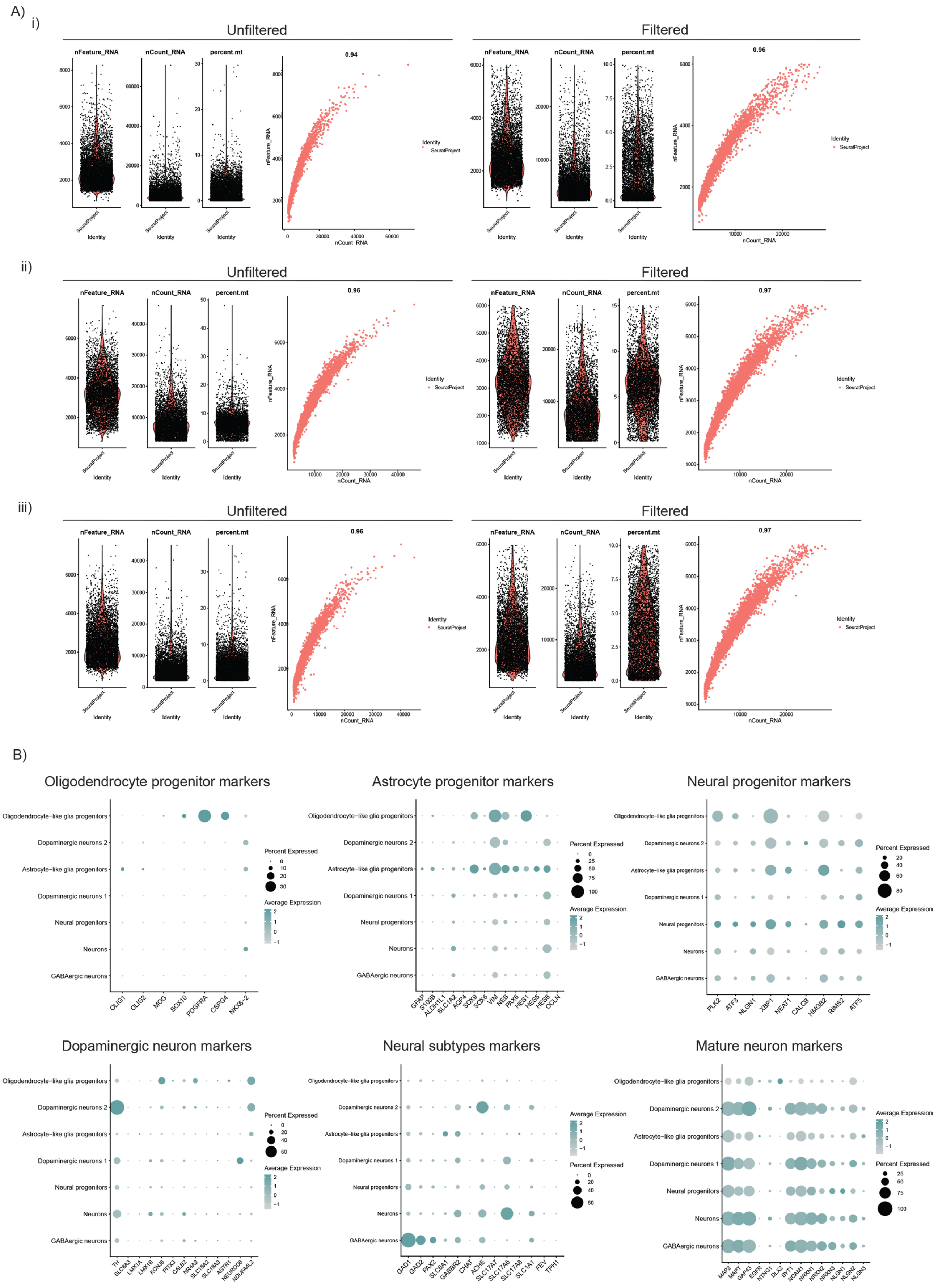


Supplementary Figure 1

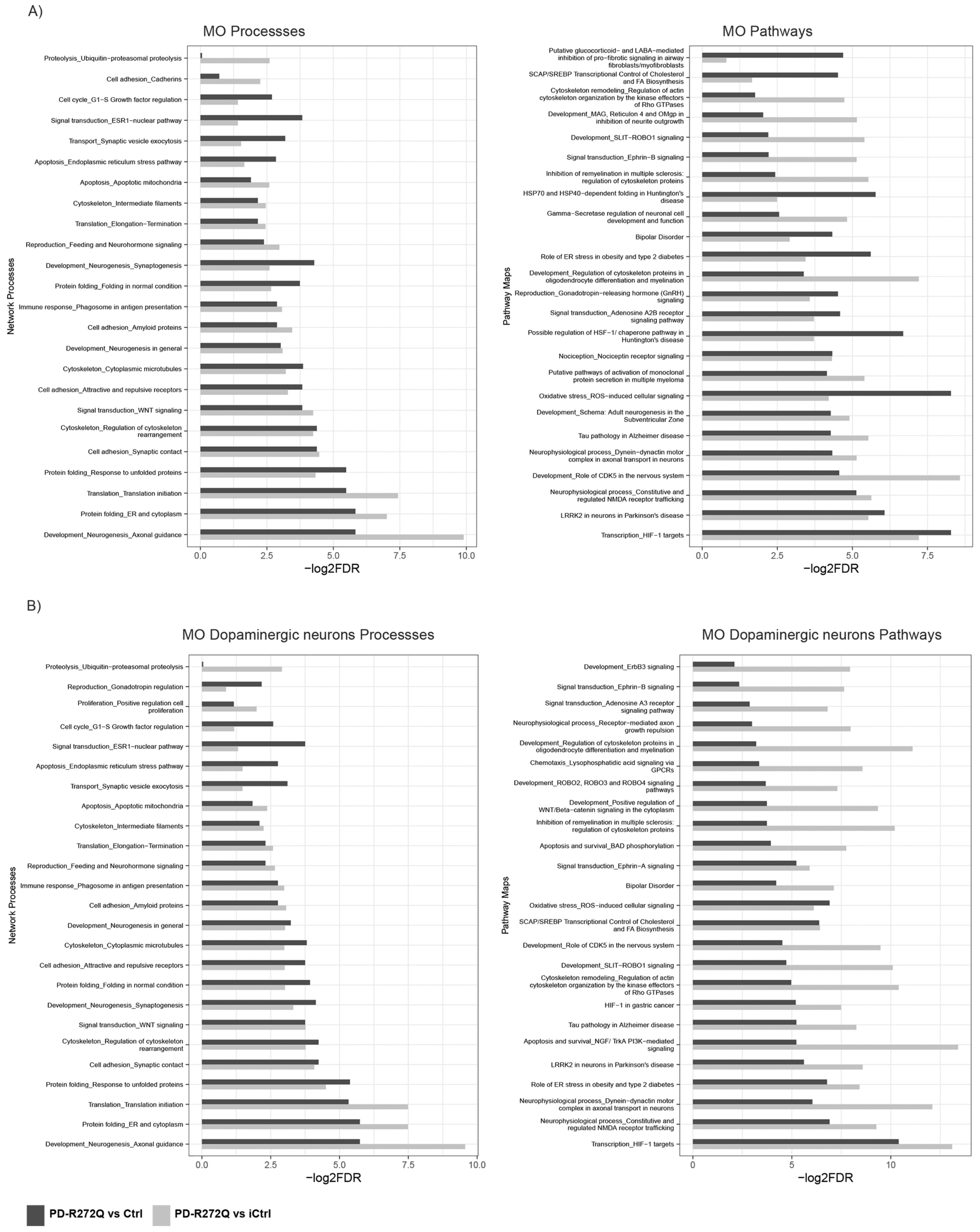


Supplementary Figure 2

Dopaminergic neuron markers

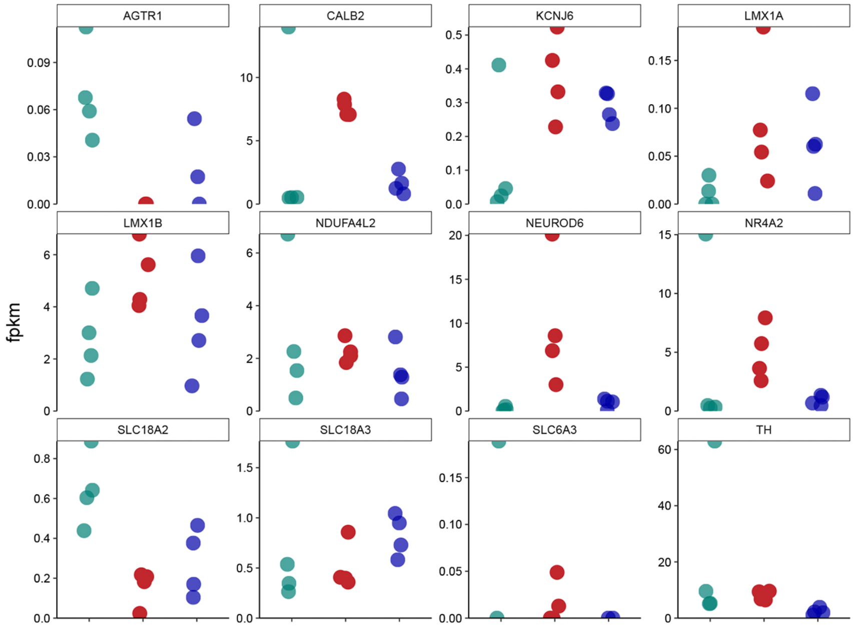


Glia markers

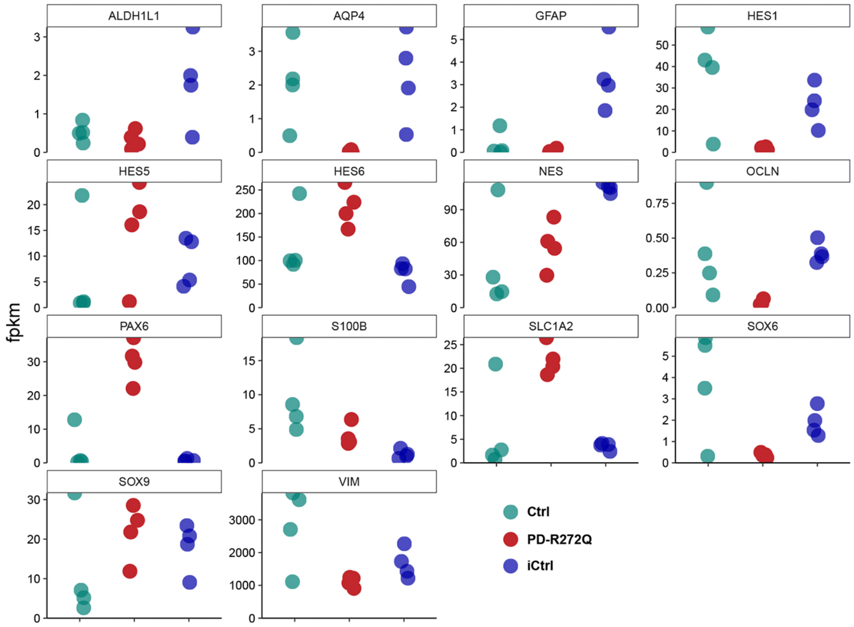


Supplementary Figure 3

MO deregulated genes in the *Oxidative Stress_ROS-induced cellular signaling* pathway

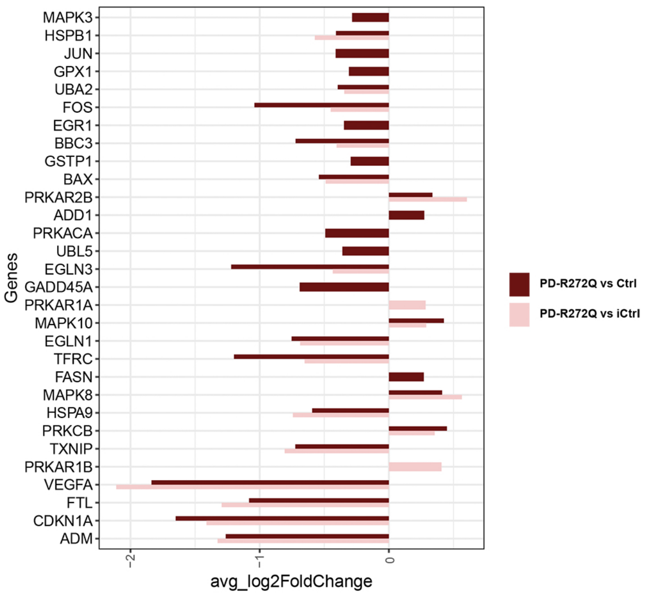


Supplementary Figure 4

A)

Midbrain organoids *RHOT1* expression levels

B)

Dopaminergic neurons *RHOT1* expression levels

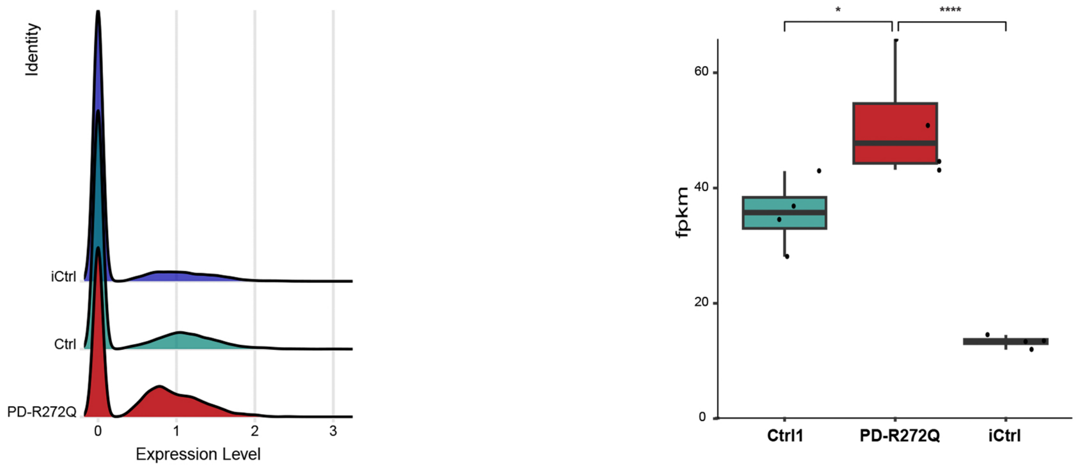


Supplementary Figure 5

Genes in “*Apoptosis_Apoptotic mitochondria*” MO dopaminergic neuron clusters deregulated process

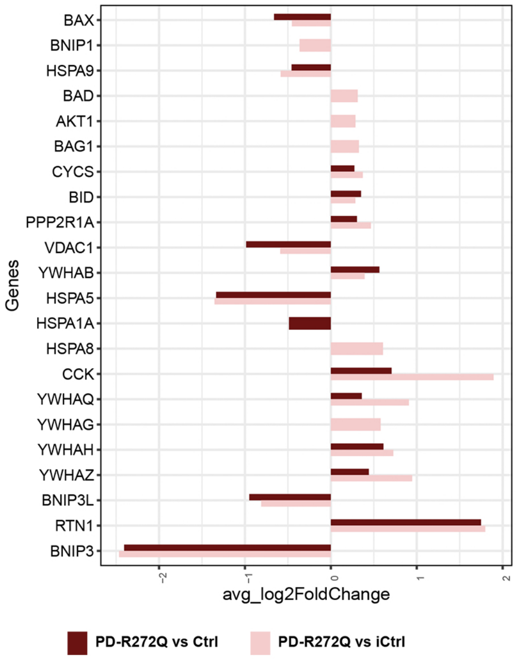


Genes in “*Apoptosis and survival_NGF/TrkA PI3K−mediated signaling*” MO dopaminergic neuron clusters deregulated pathway

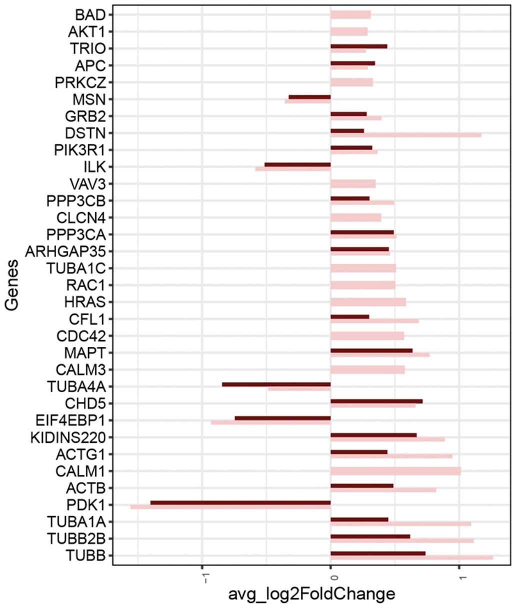


Supplementary Figure 6

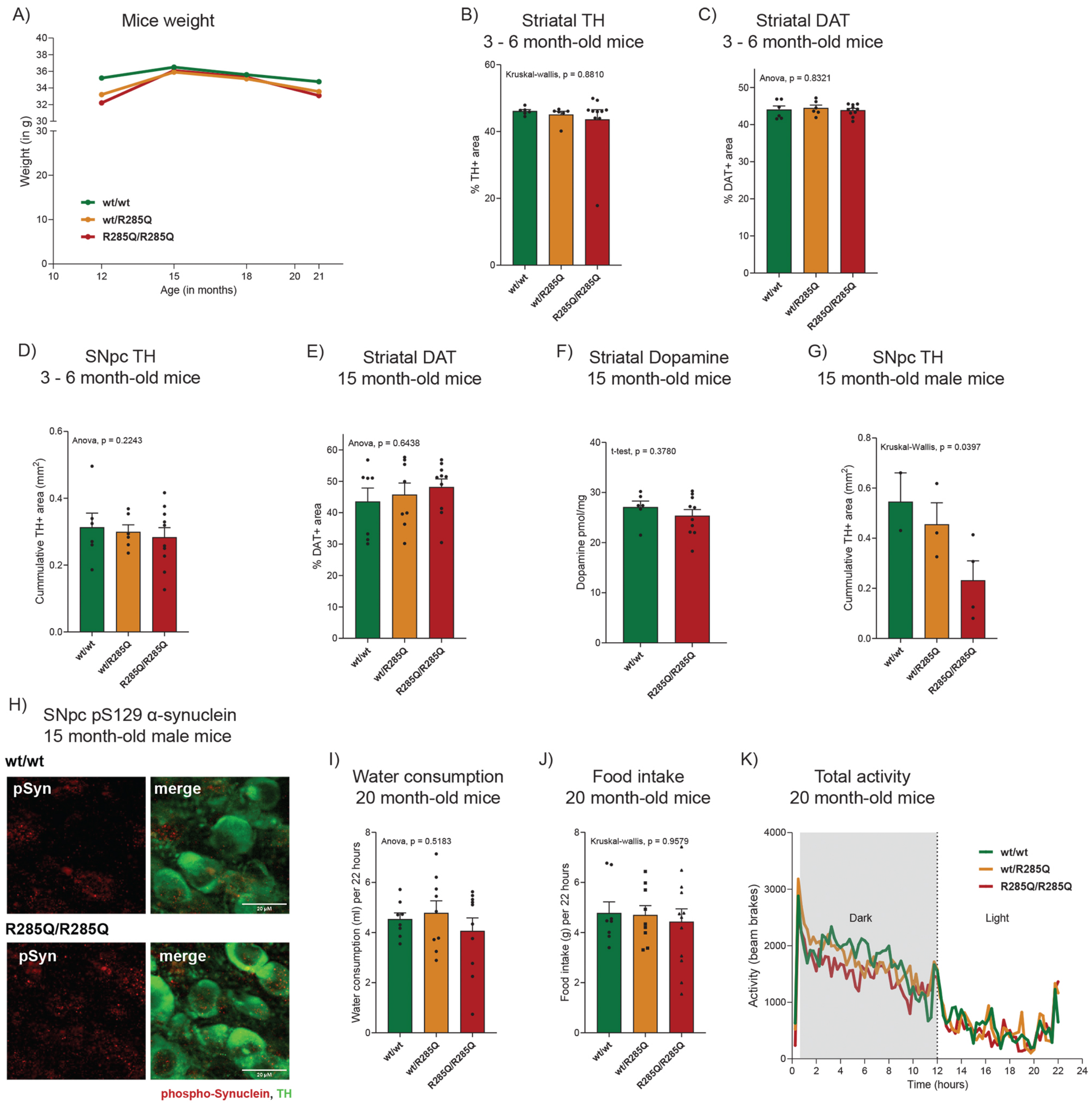


Supplementary Figure 7
