## Supplementary Material for "Parkinson’s disease-related Miro1 mutation induces mitochondrial dysfunction and loss of dopaminergic neurons *in vitro* and *in vivo*"

### Supplementary Material 1:

### Midbrain organoids metabolomics:

#### . Detailed LC-MS setting for assessing polar intracellular midbrain organoids metabolites:

*Analytical column:* SeQuant® ZIC-pHILIC 5µm polymer 150 x 2.1 mm

*Guard column:* SeQuant® ZIC-pHILIC Guard 20 x 2.1 mm

*Mobile phase A:* 20 mmol/L ammonium acetate in $H2O$ (pH 9.2, + 5 µM MA)

*Mobile phase B:* ACN (pH unadjusted, 5 µM MA)

*Gradient:*

| Time (min) | %B | Flow rate (µL/min) |
|---|---|---|
| 0.00 | 80 | 200 |
| 3.00 | 80 | 200 |
| 18.00 | 20 | 200 |
| 19.00 | 80 | 200 |
| 24.5 | 80 | 200 |
| 25.5 | 80 | 400 |
| 29.5 | 80 | 400 |
| 30.00 | 80 | 200 |

*Flow rate:* 0.20-0.40 mL/min

*Column temp.:* 45 °C

*Injection volume:* 5 µL

*Autosampler temp.:* 4 °C

**MS parameters**

*Instrument:* Exploris 240

*Ion mode:* Polarity Switching (positive/negative ESI in Full scan)

**ESI Source**

*Sheath gas flow rate*: 35

*Aux gas flow rate:* 7

*Sweep gas flow rate*: 0

*Spray voltage kV*: 3.5

*Capillary temp. (°C)*: 400

*S-lens RF level:* 70.0

*Aux gas heater temp. (°C):* 275

**Scan parameters Full MS**

*MS1 Scan range:* 75 – 1000 m/z

*Polarity:* POS/NEG

Spray voltage of 3 kV in both positive and negative mode

*MS1 Resolution:* 60,000

*MS1 AGC target:* ¨standard”

*MS1 Maximum injection time:* 100 ms

*MS1 Scan range:* 75 – 1000 m/z

*Polarity:* POS/NEG

Spray voltage of 3 kV in both positive and negative mode

*MS2 Resolution:* 30,000

*MS2 AGC target:* ¨standard”

*MS2 Maximum injection time:* “auto”

*MS2 isolation window:* 0.4 m/z

*Collision energy:* 30 V

*Loop count:* 3

*Dynamic exclusion time:* 3 s with exclusion after 1 acquisition.

**Supplementary Material 2:**

**Extracellular metabolomics on dopaminergic neurons:**

Masses used for quantification and qualification of the derivatized target analytes (dwell times between 20 and 70 ms).

| Analyte Name | Quantification Ions (m/z) | Qualification Ion I (m/z) | Qualification Ion II (m/z) |
|---|---|---|---|
| **Pyruvic acid 1MEOX 1TMS** | 174-179 | 158.1 | 189.1 |
| **Lactic acid 2TMS** | 219.1-224.1 | 190.1 | 117.1 |
| **Alanine 2TMS** | 218.1-223.1 | 190.1 | 116.1 |
| **Valine 2TMS** | 144.1 | 218.1 | 246.2 |
| **Urea 2TMS** | 189.1 | 103.1 | 171.1 |
| **Leucine 2TMS** | 158.1 | 218.1 | 232.2 |
| **Isoleucine 2TMS** | 158.1 | 218.1 | 232.2 |
| **Glycine 3TMS** | 276.1-280.1 | 248.1 | 174 |
| **Serine 3TMS** | 306.1-311.1 | 218.1 | 204.1 |
| **Threonine 3TMS** | 218.1 | 291.2 | 320.2 |
| **IS Pentanedioic acid-D6 2TMS** | 267.1 | 163.1 | 239.1 |
| **Methionine 2TMS** | 176.1 | 250.1 | 293.1 |
| **Glutamic acid 3TMS** | 363.2-371.2 | 246.1 | 348.2 |
| **Phenylalanine 2TMS** | 192.1 | 218.1 | 266.1 |
| **Asparagine 3TMS** | 231.1-236.1 | 132.1 | 348.1 |
| **IS [UL-13C5]-Ribitol 5TMS** | 220.1 | 310.2 | 323.2 |
| **Glutamine 3TMS** | 347.2-354.2 | 245.1 | 156.1 |
| **Fructose 1MeOX 5TMS** | 307.2, 310.2 | 217.1, 220.1 | |
| **Glucose 1MEOX 5TMS** | 319.2, 323.2 | 217.1, 220.1 | |
| **Lysine 4TMS** | 174.1 | 317.2 | 434.3 |
| **Tyrosine 3TMS** | 218.1 | 179.1 | 280.2 |
| **Inositol 6TMS** | 305.1 | 318.1 | 507.3 |
| **Tryptophan 2TMS** | 218.1-223.1 | 130 | 348.2 |
| **Alanyl-glutamine 3TMS** | 418.2 | 347.2 | 216.1 |
| **Cystine 4TMS** | 218.1-223.1 | 297.1 | 411.2 |
